# A rational blueprint for the design of chemically-controlled protein switches

**DOI:** 10.1101/2021.01.21.427547

**Authors:** Sailan Shui, Pablo Gainza, Leo Scheller, Che Yang, Yoichi Kurumida, Stéphane Rosset, Sandrine Georgeon, Bruno E. Correia

## Abstract

Small-molecule responsive protein switches are crucial components to control synthetic cellular activities. However, the repertoire of small-molecule protein switches is insufficient for many applications, including those in the translational spaces, where properties such as safety, immunogenicity, drug half-life, and drug side-effects are critical. Here, we present a computational protein design strategy to repurpose drug-inhibited protein-protein interactions as OFF- and ON-switches. The designed binders and drug-receptors form chemically-disruptable heterodimers (CDH) which dissociate in the presence of small molecules. To design ON-switches, we converted the CDHs into a multi-domain architecture which we refer to as *activation by inhibitor release* switches (AIR) that incorporate a rationally designed drug-insensitive receptor protein. CDHs and AIRs showed excellent performance as drug responsive switches to control combinations of synthetic circuits in mammalian cells. This approach effectively expands the chemical space and logic responses in living cells and provides a blueprint to develop new ON- and OFF-switches for basic and translational applications.

## Introduction

Synthetic biology has enabled important developments on the understanding of fundamental aspects in biology as well as in next-generation cell-based therapies^1–3^. In synthetic biology, many strategies have been pursued to control the timing, localization, specificity and strength of transgene expression or signaling events by equipping cells with sophisticated genetic circuits governed by small-molecule controlled protein switches. Typically, protein switches function are triggered by a small molecule to control the assembly or disassembly of two protein subunits^1^. One of the most widespread switch systems is the rapamycin-controlled chemically induced dimers (CIDs), FKBP:FRB which has been used in a wide variety of applications, including to control chimeric antigen receptor T (CAR-T) cell activities as safety switches^4^.

However, the large majority of chemically-controlled protein switches have limitations, particularly in translational applications, due to drug toxicity/side effects^5,6^ and unfavorable pharmacokinetics^7^, or concerns that non-human protein components could raise an immunogenic response^8–10^. Thus, an important requirement to enhance the breadth and scope of synthetic biology applications is to expand the universe of protein-based switches and consequently of the chemical space used to control engineered cellular activities.

Beyond naturally-sourced protein switches^1^, several methods have recently been proposed to expand the panel of available protein switches. Specifically for CIDs, Hill and colleagues used an *in vitro* evolution-based approach to engineer antibodies that engage with Bcl-XL only in the presence of a small-molecule drug^11^, and showed that these switches were active in cellular applications. Foight and colleagues used libraries of computationally designed mutants of a previously reported *de novo* protein scaffold to interact with a viral protease only in its drug-bound state^12^. These were also shown to regulate cellular activities in vivo, but the crystal structures of the designs evidenced substantial differences to the predicted binding modes. Remarkable computational design work was performed by Glasgow and colleagues, where a CID was rationally designed by transplanting the binding sites of a ligand to an existing protein dimer^13^. However, the precise design of key interaction residues to mediate small molecule interactions and control CIDs remain an extremely challenging computational design problem.

All these approaches focus on chemically induced dimerization systems^1,11–13^, yet chemical disruption systems also have important applications in synthetic biology and remain much less explored^14,15^. Therefore, we devised a strategy to design chemically-controlled switches by repurposing protein components and small molecules involved in the inhibition of protein-protein interactions (PPI)^16^. Multiple PPI inhibitors have been in clinical development over the past few years and some are approved for clinical use^16,17^, which make them attractive molecules for synthetic biology applications in the translational domain.

Previously, we reported the design of chemically disruptable heterodimers (CDH) from known protein: peptide-motif complexes, by transferring the peptide motif from the disordered binding partner to globular proteins and using the known PPI inhibitor as a chemical disruptor^18^. Specifically, our CDH was based on the Bcl-XL:BIM-BH3 complex, where the BIM-BH3 interaction motif was transplanted to the Lead Design 3 (LD3) using computational design, resulting in a Bcl-XL binder with 1000-fold times higher affinity than the wildtype disordered BIM-BH3 peptide-motif. Biochemically, we showed that Bcl-XL:LD3 complex (CDH-1) is disrupted by the drug A-1155463 (Drug-1)^19^, and can be used as an OFF switch for CAR-T cell activity in a dose-dependent, dynamic, and reversible manner *in vivo*^18^.

Here, we expand our computationally-driven approach to design novel chemically controlled protein switches using distinct small-molecule drugs. First, we assembled a novel CDH based on the Bcl2:LD3 protein complex (CDH-2) disrupted by the clinically-approved drug Venetoclax (Drug-2)^21,22^. A novel CDH-3 was designed based on the interaction of the mdm2 protein and the p53 peptide motif (disrupted by Drug-3^23^) (Fig. 1a). The three CDHs (CDH-1-3) were used to regulate cellular responses, which included transcriptional gene expression^24^ and synthetic cell surface receptor signaling based on the GEMS platform^25^. In order to investigate the impact of biochemical parameters of the CDHs on drug disruptor dosage and their cellular activity, we designed a suite of CDHs with tailored affinity ranges from low picomolar to mid nanomolar and tested their effect in regulating cellular processes (Fig. 1b). Next, we devised a novel strategy to create ON switches (i.e. protein pairs where a small-molecule drug promotes heterodimerization) by repurposing the CDHs. These new switches, referred to as *activation by inhibitor release switch* (AIR), have a “split” architecture with two protein chains where one of the chains contains the two domains of a CDH genetically fused, and the second chain contains a rationally-designed drug insensitive receptor that retains the ability to bind to the computationally designed binder (e.g. LD3). The drug disruptor releases the designed binder from the fused CDH exposing the interaction site of the binder and consequently enabling the dimerization with the drug insensitive receptor (Fig. 1c). Notably, as the array of the clinically-validated PPI inhibitors grows^16^, our AIR strategy could be used to accelerate the development of ON-switches to control activities in engineered cells, which remains an important need in synthetic biology (Fig. 1d). Finally, we engineered mammalian cells equipped with multi-input/output control modes which showcased the broad applicability of the designed protein switches into more sophisticated control modes broadening the potential scope of applications (Fig. 1e).

**Figure 1.**
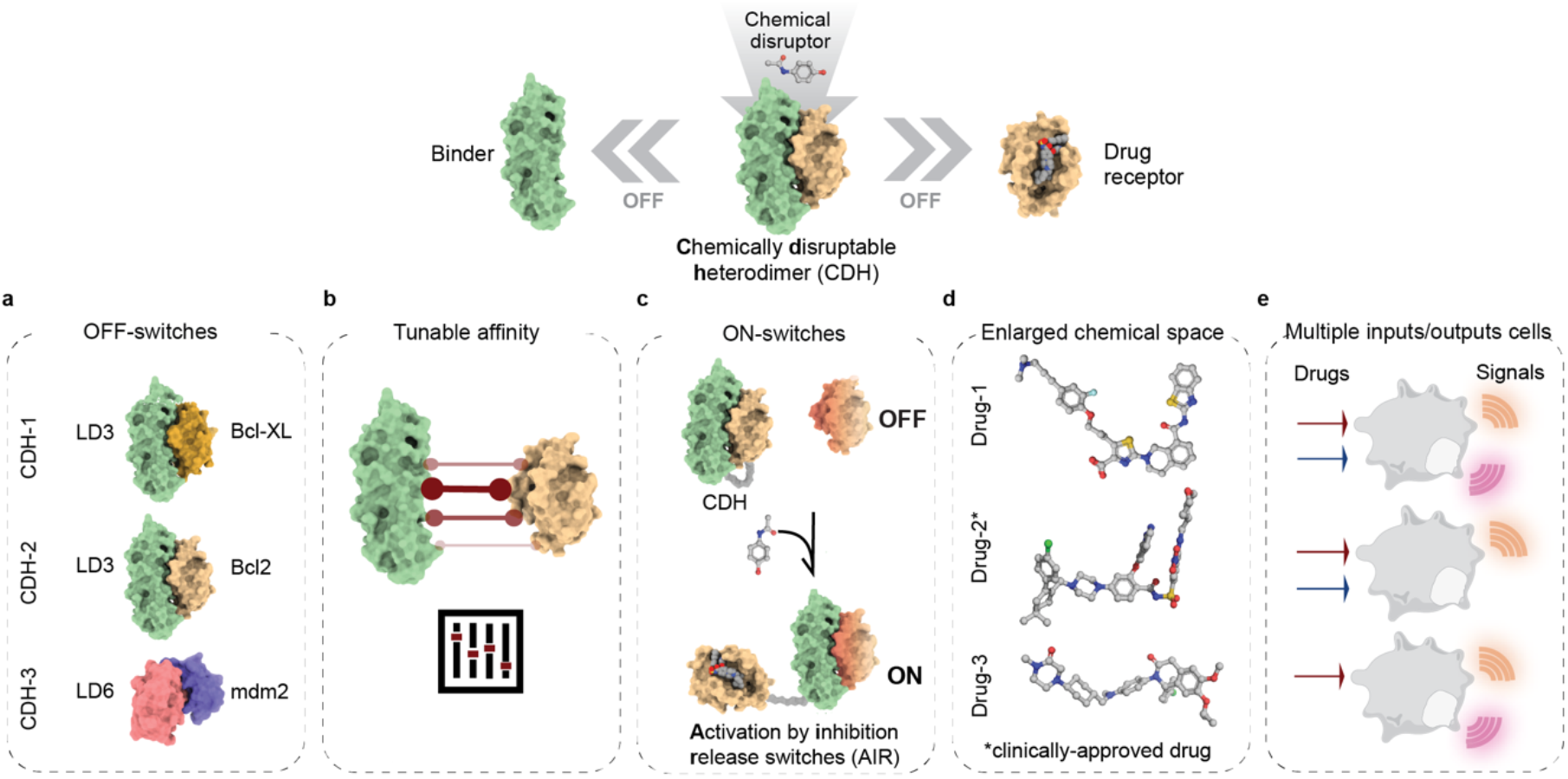
Overview of structure-based design strategies employed to create novel smallmolecule responsive switches. These design strategies include: I) binding site transplantation to create new CDHs **(a)** that can be used to control cellular activities; II) Interface mutations for affinity tuning **(b)** to alter drug concentrations that interfere with assembly of the complexes; III) Multistate design to create drug insensitive receptors that are embedded in multi-domain architectures that dimerize upon drug exposure **(c)**. This protein switch toolbox enlarges the chemical space **(d)** and can be used for the engineering of cells which sense several chemical cues and output a variety of signals **(e)**.

Altogether, we provide a rational blueprint that leverages structure-based and computational design approaches to design novel small-molecule switches for use across basic and translational synthetic biology applications.

## Design of CDHs controlled by clinically relevant drugs

Recently, we described the design of CDH-1 composed of the Bcl-XL:LD3 complex which dissociates in the presence of the Bcl-XL-specific inhibitor A-1155463 (Drug-1). However, Drug-1 is not a clinically-approved drug, which limits the potential translational applications of the CDH-1. We found that LD3 also binds to Bcl2 (Fig. 2a), a protein from the Bcl2 family^26,27^ closely related to Bcl-XL, with a dissociation constant (K_d_) of 0.8 nM as determined by surface plasmon resonance (SPR) (Fig. 2b). Bcl2 is the target of Venetoclax^28^ (Drug-2) (Supplementary Fig. 1a,b), a BH3 chemical mimetic that blocks the anti-apoptotic Bcl2 protein and is clinically-approved for chronic lymphocytic leukemia (CLL) treatment^22^. We therefore assembled CDH-2, where we exchanged the Bcl-XL component with Bcl2 (Fig. 2a). Drug-2 effectively disrupts the CDH-2 heterodimer *in vitro* both in an SPR drug competition assay (Fig. 2c, Supp. Fig. 1c) (IC_50_ = 67.5 nM), as well as by the elution profiles of size-exclusion chromatography coupled with multi-angle light scattering (SEC-MALS) (Supp. Fig. 1d).

**Figure 2:**
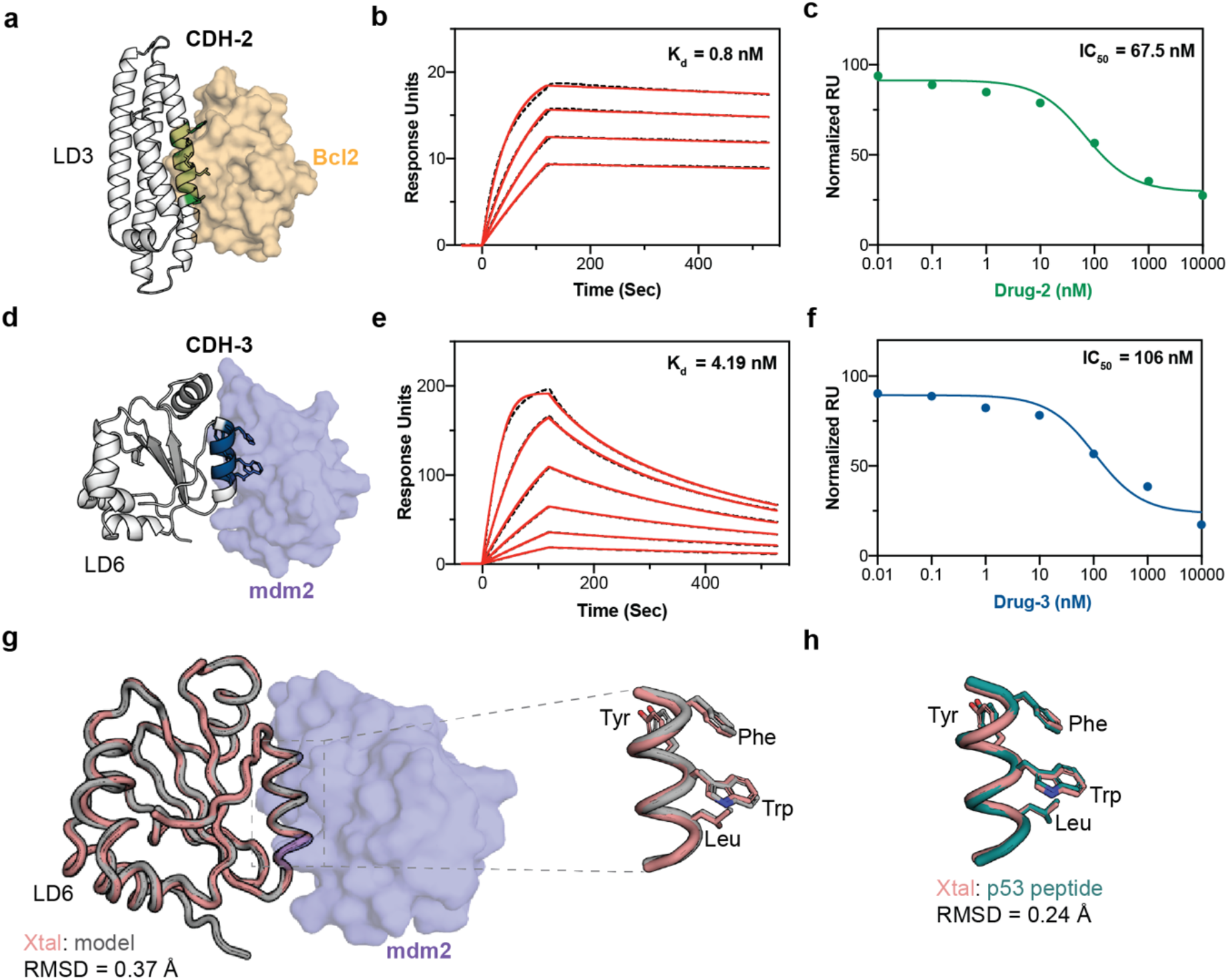
Biochemical and structural characterization of novel CDHs. **a)** Structural representation of CDH-2 composed of Bcl2 (beige surface) and LD3 (white cartoon) with the interfacial segment colored in green, the hotspot residues transplanted from the BH3 motif are shown in sticks. **b)** SPR measurements of CDH-2 binding affinity. The dissociation constant determined for the interaction of Bcl2 with LD3 is 796 pM. **c)** SPR drug competition assay determined the apparent IC_50_ of CDH-2 with Drug-2 around 67.5 nM. **d)** Structural representation of CDH-3 composed of the mdm2 (purple surface):LD6 (white cartoon) complex with the interfacial segment colored in blue, the hotspot residues transplanted from the p53 are motif shown in sticks. **e)** SPR measurements of CDH-3 binding affinity. The dissociation constant determined for the interaction of mdm2 with LD6 is 4.19 nM. **f)** SPR drug competition assay determined the apparent IC_50_ of CDH-3 with Drug-3 around 106 nM. **g)** The crystal structure of the CDH-3 consisting of the complex LD6 (pink tube) with mdm2 (purple surface) was in close agreement with the computational model of LD6 (gray tube) with complex with mdm2 (not shown). **h)** Transplanted hotspot residues in LD6 in sticks (pink) aligned with p53 peptide residues (teal) in a RMSD of 0.24 Å.

To further expand the panel of CDHs and the chemical space of small-molecule disruptors, we designed a novel CDH-3 (Fig. 2d), based on the interaction between p53 and mdm2 as the starting point. Over the past two decades, numerous compounds have been developed to inhibit this interaction and several candidates have been tested in clinical trials^29,30^. Particularly, the dihydroisoquinoline derivative, NVP-CGM097 (Drug-3) (Supp. Fig. 2a,b), is currently undergoing phase 1 clinical trials^16,23,31^. Leveraging the same computational design approach described for the CDH-1^18^, we utilized the structural information of the mdm2:p53 stapled peptide complex (PDB ID: 5afg)^32^ and searched for proteins that could accommodate the p53 helical motif. We performed a structural search over approximately 11,000 putative protein scaffolds and after the design stage we selected for experimental characterization three designs which we refer to as LD4, LD5 and LD6 (Supp. Fig. 2c-f, Fig. 1e). From these three designs, LD6 a design based on a thioredoxin protein from mouse^33^, showed the best biochemical behavior being monomeric in solution (Sup. Fig. 3a) and a melting temperature of 62 °C (Supp. Fig. 3b). LD6 bound to mdm2 with a K_d_ of 4 nM (Fig. 2e) as determined by SPR, showing a higher binding affinity than those reported for the wild-type p53_17-29_ peptide (820±60 nM) or the stapled p53 peptide which we used as the input peptide motif (12±3 nM)^32^. Next, we tested if Drug-3 promoted the dissociation of the mdm2:LD6 complex by SPR and determined an IC_50_ of 106 nM (Fig. 2f, Supp Fig. 3c). These results were also confirmed using SEC-MALS where the drug dissociated the complex into two monomers (Supp Fig. 3d).

To evaluate the structural accuracy of the designed CDH-3, we solved the crystal structure of the mdm2:LD6 complex at 2.9 Å resolution by X-ray crystallography. The structure of the complex closely matched our computational model, with backbone RMSDs of 0.37 Å over the overall structure (Fig. 2g) and 0.24 Å over the p53 motif region (Fig. 2g, Supp. Fig. 4). The conformation of key residues observed in the native interface were closely mimicked in the crystal structure of LD6 in complex with mdm2. Overall, our results show that we can robustly use computational design to create protein modules which the assembly state is controlled by the presence of small-molecules.

## CDHs function as intra and extra-cellular OFF-switches

To test whether the designed CDHs could be useful for distinct synthetic biology applications, we generated OFF-switches for two cellular systems: transcriptional gene regulation and control of endogenous signaling pathways (Fig. 3a, d). To establish reliable drug concentration ranges for our experiments, we first assessed drug toxicities and impact on the expression of reporter proteins in HEK293T cells, Drugs1-3 were well tolerated up to 10 μM with no detectable changes in the expression of reporter proteins (Supp. Fig. 5a, b).

**Figure 3.**
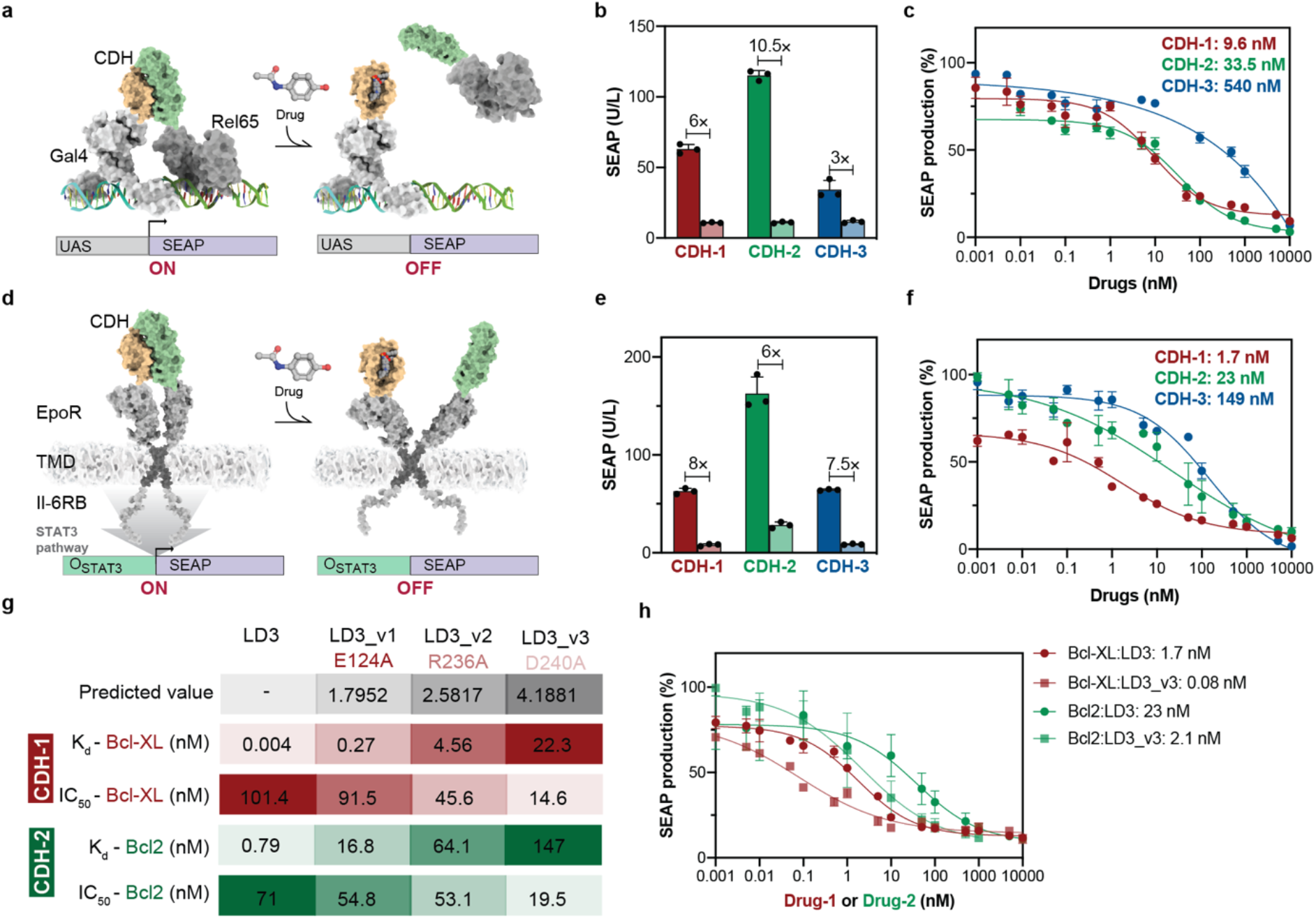
Exploration of distinct cellular activities, localization and affinities of the CDH switches. **a)** Schematic representation of the CDHs utilized to control transcription regulation with the GAL4/UAS system (CDH-TFs). **b)** CDH-TFs fold-change activity between drug treated (1 μM) versus untreated (DMSO), showing the quantification of SEAP expression after 24 h drug treatment. **c)** Drug dosedependent responses of CDH-TFs quantified by SEAP expression after 24 h drug treatment. **d)** Schematic representation of the CDHs utilized to regulate surface signaling receptors (CDH-GEMS) in engineered cells. **e)** CDH-GEMS fold-change activity between drug (1 μM) versus no drug treatment (DMSO), showing the quantification of SEAP expression after 24 h. **f)** Drug dose-dependent responses of CDH-TFs quantified by SEAP expression after 24 h. **g)** Summary of the LD3 mutants including computationally predicted decreases in affinity and experimental measurements using SPR. Values shown are the predicted interaction energy of LD3 mutants (ΔΔG in Rosetta energy units), the binding affinities with Bcl-XL and Bcl2 and IC_50_s of CDH dissociation also for Bcl-XL and Bcl2 with Drug-1 and Drug-2, respectively. **h)** Drug dose-dependent responses in engineered cells determined for the CDH-(1-2)-GEMS with LD3 and LD3_v3. The drug receptor utilized were Bcl-XL (CDH-1) in red and Bcl2 (CDH-2) in green, CDHs with original LD3 were in circle symbol and LD3_v3 in square symbol with Bcl-XL and Bcl2. **b, e, c, f, h**) Each data point represents the mean ± s.d. of three replicates and the IC_50_s were calculated using four-parameter nonlinear regression.

To regulate transgene activation events by small molecule inputs we used the CDHs with a split transcription factor design (CDH-TFs) based on the well-characterized Gal4/UAS system^24^ (Fig. 3a).

In brief, the CDH mediated dimerization of the Gal4 DNA binding domain and the Rel65 activation domain trigger the transcription of the reporter protein (secreted alkaline phosphatase (SEAP)). If the CDH switch is functional, the drugs should dissociate the engineered CDH-TFs and diminish SEAP expression. Upon drug treatment, all CDH-(1-3)-TFs were responsive to their respective drugs with dynamic response ranges between 3 and 10-fold (untreated vs 1 μM Drug) (Fig. 3b). The CDH-TFs showed dose-dependent responses with IC_50_s of 9.6, 33.5 and 540 nM for CDH-1-TF, CDH-2-TF, and CDH-3-TF, respectively (Fig. 3c). The CDH-TF responses were drug-specific, as each CDH-TF only responded to their respective drug (Supp. Fig. 6a). Finally, we explored the dynamic behavior of the designed components by subjecting the CDH-TF transfected cells to intermittent drug treatments. The CDH-TFs showed a significant decrease upon drug treatment and subsequent recovery of SEAP expression upon drug withdrawal (Supp. Fig. 6b-c), exhibiting a reversible behavior which is central for many synthetic biology applications. Together, these results show that our extended panel of CDHs can be used as intracellular switches and may likely be adaptable to control several cellular activities in engineered cells that are dimerization/co-localization dependent.

To test modularity and performance robustness of the CDHs in different molecular contexts, we inserted these modules in the generalizable extracellular molecule sensor (GEMS) receptor signaling pathway platform^25^ and built CDH-GEMS. In these constructs the CDHs are located extracellularly and function as ectodomain switches, fused to the backbone of the erythropoietin receptor, and an intracellular interleukin 6 receptor domain which activates the JAK/STAT pathway (Fig. 3d). The CDH-GEMS trigger the activation of the JAK/STAT pathway by the dimerization of their subunits. Thus, the drugs are expected to split the CDHs inactivating the signaling pathway as assessed by the expression of the reporter protein SEAP. All three CDH-GEMS were responsive to their respective drugs dynamic response range varied from approximately 6-fold (CDH-2) to 8-fold (CDH-1,3) (Fig. 3e). They also showed a dose-dependent response with IC_50_s of 1.7, 23 and 149 nM respectively for CDH(1-3)-GEMS with their cognate drugs (Fig. 3f). The CDH-GEMS only function when both chains are transfected (Supp. Fig. 6d) and are specifically inhibited by their corresponding drugs (Supp. Fig. 6e). The CDH-GEMS were also reversible upon intermittent drug treatment (Supp. Fig. 6f-g). Despite that comparisons between CDH-TFs and CDH-GEMS are not straightforward due to differences in the reporter signaling pathways, we observe a general trend for the CDHs to present lower IC_50_s when used as extracellular modules which is consistent with overcoming drug permeability issues in cells.

Our design process optimized the CDHs for the strength of their interaction, achieving affinities as tight as 4 pM (CDH-1). However, it is unclear how strong the interaction for a specific application should be, and, indeed, many intracellular pathways effectively function with weak protein interaction affinities^34^. We hypothesized that the sensitivity of a CDH could be tuned towards lower drug demand, by weakening the interaction of the two CDH components. We performed computational alanine scanning on the designed binder LD3 to generate lower-affinity binders and selected three mutants (LD3_v1-v3) (Supp. Fig. 7a-c) which were predicted to decrease the interaction energy (ΔΔG) to different extents (Fig. 3g). LD3 affinity to Bcl-XL measured by SPR was approximately 4 pM and the weakest design (LD3_v3) showed a binding affinity of 22.3 nM, a decrease of more than 5,000-fold (Fig. 3g, Supp. Fig. 7d-f, j); in vitro IC_50_s also lowered from 101 nM to 14.6 nM, a 7-fold reduction. Next, we tested whether this decrease in affinity also translated into an increased drug sensitivity in CDH-GEMS engineered cells. We observed a 21-fold reduction in IC_50_s of the weakest binder LD3_v3 in Bcl-XL-based CDH-GEMS when compared to the original LD3, showing that we successfully decreased the amount of drug necessary to control the CDH in cells (Supp. Fig. 7j, Fig. 3h). We also tested the panel of LD3 mutants in the context of the CDH-2 and observed similar trends (Supp. Fig. 7g-i, k; Fig. 3g, i), using a clinically approved drug.

While the overall trends were maintained, the magnitude of the reductions in binding affinities of the CDHs did not directly translate to reductions in cellular activity assays, which may reflect a number of factors associated to the complexity of performing measurements in living cells. In summary, the reduced binding affinity increased the sensitivity of the CDHs towards their small-molecule drugs. This property enlarges the panel of molecular switches that can function under reduced drug dosages and could avoid potential toxicities while remain effective at controlling engineered cell activities.

## Computational design of drug-controlled dimerizing ON switchess

CDH components are inherently well suited to trigger OFF outputs, which are highly desirable in some settings, as we previously demonstrated by its integration in CAR-T cells^18^. However, switches that can induce protein colocalization are naturally better suited to obtain ON outputs. We propose that monomerization inducing components (e.g. CDHs) can be rapidly repurposed into dimerization inducing systems (e.g. CIDs), thus diversifying the protein switches available and expanding the chemical space of CID systems to control ON outputs.

To address this challenge, we developed a new CID switch architecture, dubbed *activation by inhibitor release* (AIR) switches (Fig. 4a). The AIR architecture relies on three protein components: CDH drug receptor, CDH protein binder and a rationally designed drug-insensitive CDH receptor that retains the binding capability to the protein binder. These three components are assembled into two distinct polypeptide chains to form the AIR switches. In one chain the two elements of the CDH are fused with a flexible peptide linker forming an intramolecular binding interaction that will be disrupted in the presence of the drug, unveiling the binding site of the protein binder (Fig. 4a). In the AIR’s second chain, a drug-insensitive receptor presents the binding site to mediate the intermolecular interaction between the two chains when the drug is present. By creating this multidomain architecture, with tailored components for drug sensitivity, we aim to design protein switches that heterodimerize through an allosteric drug-binding event, effectively mediating a chemically-induced proximity mechanism. Crucial to our AIR architecture is the ability to rationally design drug insensitive receptors that can retain binding activity to the designed binders. In computational design this relates to the well-defined problem of multistate optimization where the sequence space is searched to optimize simultaneously several objective functions^35,36^. In our case, residues in the drug receptor close to the drug and away from the designed binder were selected and sampled for mutations that knocked-out drug binding (*negative* design) and maintained affinity to the protein binder (*positive* design)^37–39^ (Fig. 4a) (see Methods). We first applied this protocol to the CDH-1 (Bcl-XL:LD3), six residues in Bcl-XL (D98, R102, F105, T109, S145 and A149) were selected for multistate design (Fig. 4b), and five putative drug-insensitive Bcl-XL mutants (iBcl-XL_v1-5) were tested (Supp. Table4). iBcl-XL_v3 (R102E, F105I) and iBcl-XL_v5 (E98S, F105I) showed the greatest drug resistance (Supp. Fig. 8a) and did not dissociate from LD3 in the presence of 10 μM of Drug-1, which was sufficient to fully dissociate the wildtype Bcl-XL (Fig. 4c, Supp. Fig. 8b). The iBcl-XL_v3:LD3 retained a high affinity interaction (K_d_ = 3.8 nM) (Supp. Fig. 8c), which was however considerably lower than Bcl-XL:LD3 complex (K_d_ = 4 pM). Structural analysis of the negative design Bcl-XL:Drug-1 complex (PDB id: 4QVX)^19^ shows that F105I, shared by both designs, removes a Pi-stacking interaction with Drug-1, suggesting this mutation as the main resistance driver. To confirm the drug resistance of iBcl-XL_v3 in a cellular context, we tested the iBcl-XL_v3 paired with LD3 in the CDH-GEMS platform. In contrast to the CDH-1-GEMS, Drug-1 failed to disrupt the iBcl-XL_v3:LD3 complex at μM concentrations (Supp. Fig. 8d).

**Figure 4.**
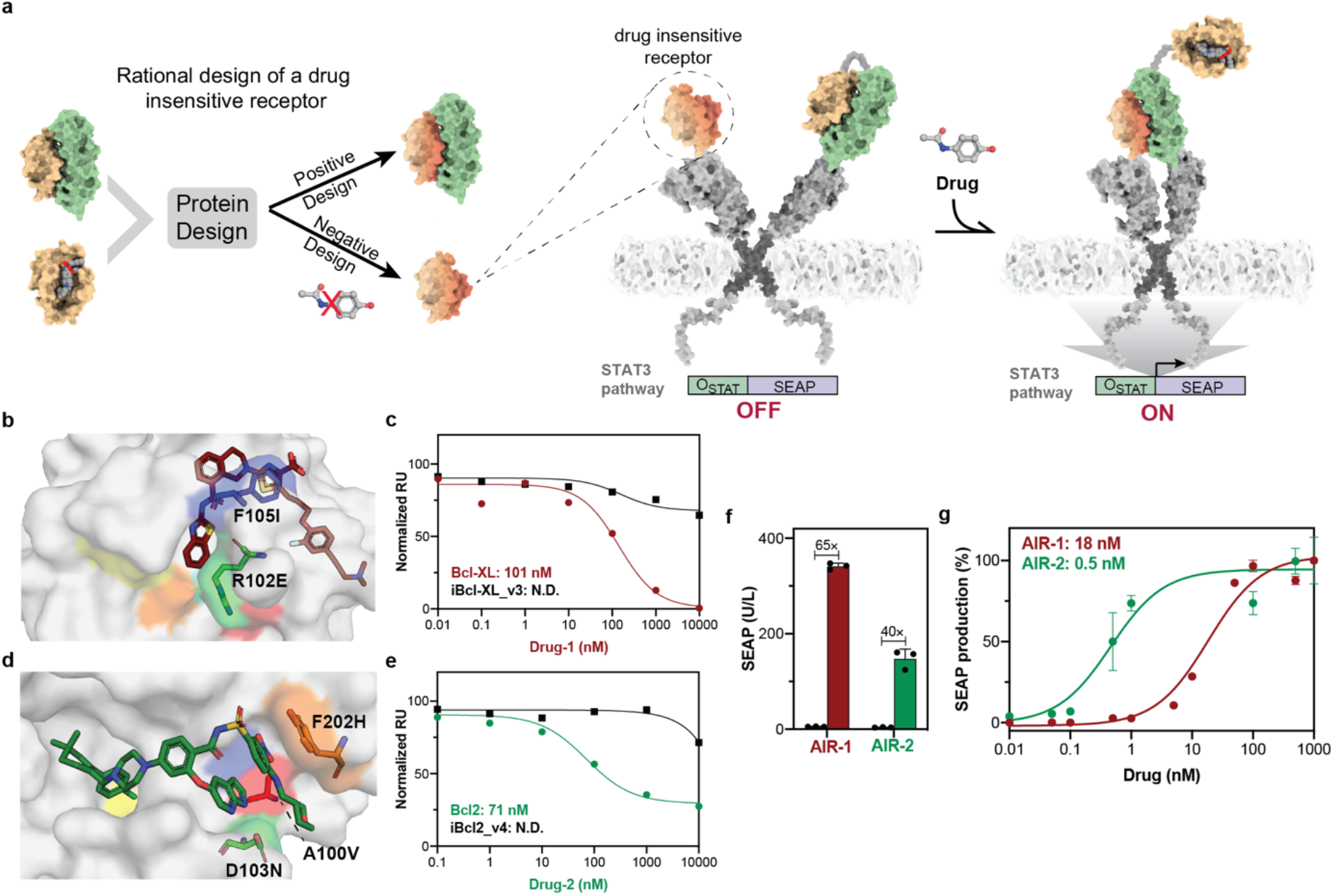
Computational design of protein components to create chemically controlled ON-switches. **a)** Computational design approach and architecture of the AIR-GEMS system. Starting from the CDH components (drug receptor (beige) and protein binder (green) we used a multistate design approach to search for drug-receptor variants that retained binding to the protein binder (positive design) and become resistant to the drug (negative design). We then assembled the AIR-GEMS, were the drug triggers the expression of SEAP by activating the JAK-STAT pathway. **b)** Structural representation of Drug-1 binding pocket in Bcl-XL. Drug binding pocket (white surface) where the mutations R102E (green sticks) and F105I (blue sticks) were performed to obtain the variant iBcl-XL_v3. Drug-1 is shown in sticks representation and colored in brown. Four other designable residues are highlighted on the surface, E98 in red, T109 in cyan, S145 in orange and A149 in yellow. **c)** Apparent IC_50_s for Drug-1 induce dissociation of Bcl-XL:LD3 and iBcl-XL_v3:LD3 determined by SPR drug competition assay. **d)** Structural representation of Drug-1 binding pocket in Bcl2. Drug binding pocket (white surface) where the mutations A100V (red sticks), D103N (green sticks) and Y201H (orange sticks) were performed to obtain the variant iBcl2_v4. Drug-2 is shown in sticks representation and colored in green. Two other designable resides are highlighted on the surface, V148 in blue, V156 in yellow. **e)** Apparent IC_50_s of Drug-2 induce dissociation of Bcl2:LD3 and iBcl2_v4:LD3 determined by SPR drug competition assay. **f)** AIR-GEMS fold-change activity between drug (1 μM) versus no drug treatment (DMSO), showing the quantification of SEAP expression after 24 h. **g)** Drug dose-dependent responses in engineered cells expressing the AIR-GEMS. Each data point represents the mean ± s.d. of three replicates and the EC_50_s were calculated using four-parameter nonlinear regression.

Next, we assembled the AIR sensors in the GEMS platform (Fig. 4a, right) to evaluate its potential as an effective ON switch. We fused the AIR components in the ectodomains of the GEMS and measured SEAP activity as a reporter for drug-triggered activation. We constructed the AIR-1-GEMS using iBcl-XL_v3 in one chain and a fused CDH-1 (LD3-GGGGSX3 linker-Bcl-XL) in the second chain. We coexpressed these constructs in HEK293T cells, and observed that the AIR-1-GEMS was responsive to Drug-1, effectively turning ON the expression of the reporter gene SEAP. Furthermore, the AIR switches showed a very sensitive drug response (EC_50_ = 18.68 ± 4.65 nM) (Fig. 4g), and a 64-fold dynamic response range (untreated vs 100 nM Drug-1 treated) (Fig. 4f).

To test whether we could design a second ON-switch controlled by a different drug, we applied the same design strategy to generate a Drug-2 resistant receptor and created the AIR-2 based on the CDH-2 components (Bcl2:LD3) which is responsive to the clinically approved drug Venetoclax (Drug-2). Similarly to the AIR-1, five residues located in the drug binding site (A100, D103, V148, V156, Y202) (Fig. 4d) were used for multistate design to create drug-insensitive Bcl2 variants (iBcl2). Four designs (Supp. Table 5) were screened on the AIR-GEMS platform and one showed activation upon addition of Drug-2 (Supp. Fig. 9a). We refer to this switch as the AIR-2-GEMS, which was composed of iBcl2_v4 (A100V, D103N, Y202H) and the fused CDH-2 (LD3-GGGGSX3 linker-Bcl2), and showed an EC_50_ of 0.5 ± 0.27 nM (Fig. 4g). Subsequently we verified at the biochemical level that it retained the interaction with LD3 (K_d_ = 9.8 μM) (Supp. Fig. 9b) and was resistant to Drug-2 both in vitro (Fig. 4e, Supp. Fig. 9c) as well as in cell-based assays (Supp. Fig. 9d).

A comparison between the two AIR-GEMS shows that AIR-2-GEMS is approximately 36-fold more sensitive to drug treatment in terms of EC_50_s, but AIR-1-GEMS shows a higher magnitude of response when comparing the resting and the fully activated states (350 U/L SEAP in AIR1-GEMS vs 160 U/L in AIR2-GEMS at 100 nM Drug) (Fig. 4f). These distinct behaviors may be due to differences in affinities of LD3 to the different drug receptors, where a lower affinity affords a more responsive receptor requiring lower drug concentrations, but the output is less sustained reaching overall lower activation levels.

These results suggest that the conversion of CDH components into the AIR architecture is an effective way to design ON switches responsive to low-doses of small-molecule drugs (including clinically approved). As the panel of clinically-tested PPI inhibitors grows, our AIR design strategy could be a general approach to expand the panel protein switches to control cellular activities.

## Multi-input multi-output control in engineered cells using orthogonal switches

An overarching goal in synthetic biology is to use living cells as bio-computing units, where orthogonal sensing components display distinct switching behaviors^40^. The need for these complex logic devices goes beyond basic applications, as shown in the CAR-T cell field, where there is a growing need to control multiple functions using orthogonal signals (e.g. suicide switches^41^, tunable activity^42^, molecule secretion^43^). As a final proof-of-concept, we engineered multi-input/multi-output cells by combining several designed switches in HEK293T cells, as a model system for further applications. In analogy with control systems in electrical engineering, we sought to implement three distinct input/output systems beyond single-input/single-output (SISO), such as: multi-inputs/single-output (MISO), single-input/multi-outputs (SIMO) and multi-inputs/multi-outputs (MIMO)^44^.

We first engineered MIMO cells to respond to two drugs with distinct outputs (Fig. 5a) by co-transfecting the CDH-3-TF with either AIR-1-GEMS (Fig. 5b) or AIR-2-GEMS (Fig. 5c). The combination of two drugs did not affect cell viability at the highest concentration of 1 μM each (Supp. Fig. 10a). The CDH-3-TF luciferase reporter expression decreased in a Drug-3 dose-dependent manner and AIR-GEMS (AIR-1 and AIR-2) showed an increase of SEAP expression dependent on their cognate drugs (Fig. 5b-c). Showing an orthogonal control of two distinct cellular activities by two drug inputs in the engineered cells.

**Figure 5.**
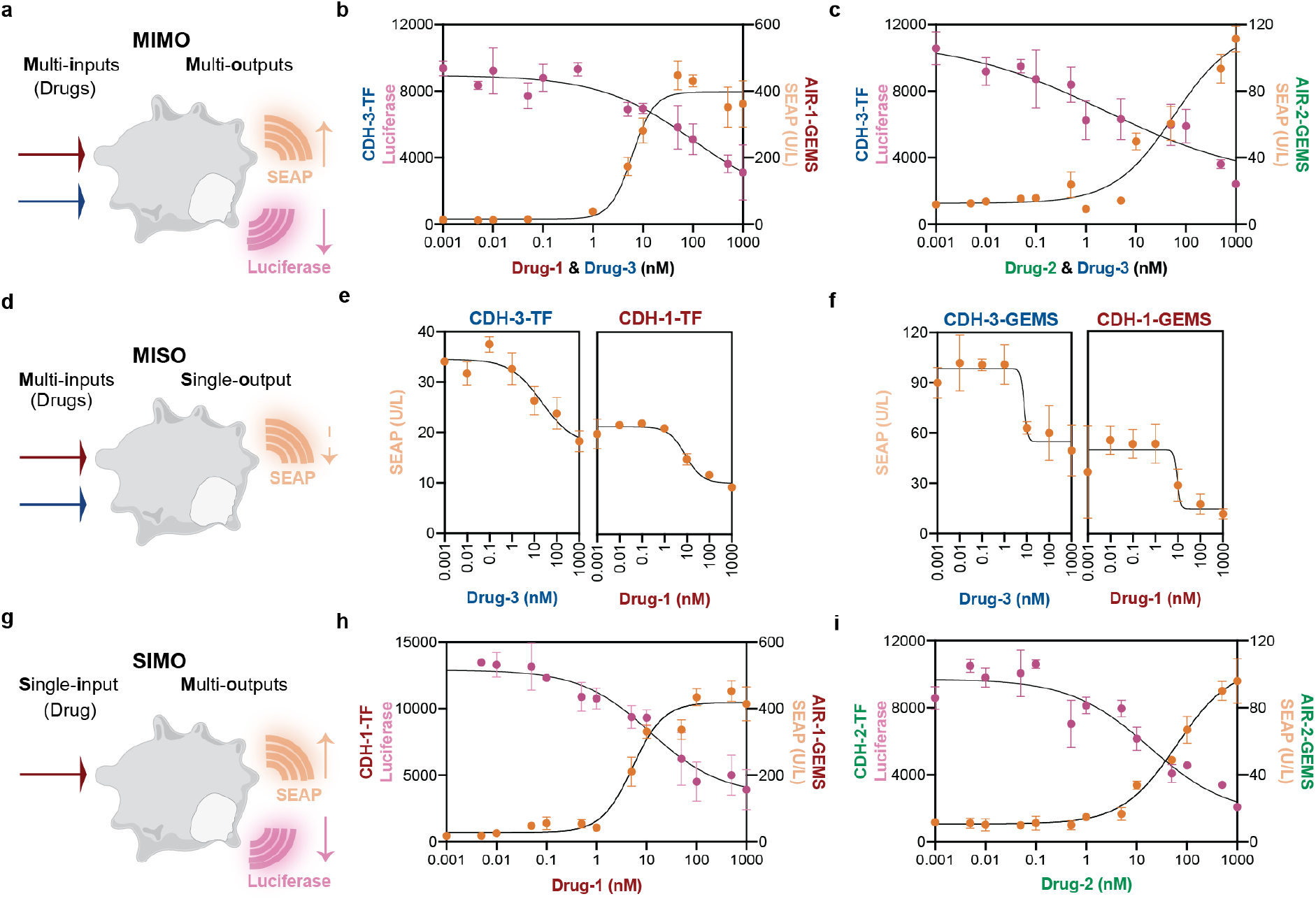
Orthogonal chemical switches enable implementation of multi-input multi-output control modes in mammalian cells. **a)** Scheme of multi-inputs multi-outputs (MIMO) designer cells where two drugs control two different outputs. **b, c)** Cells were co-transfected with AIR-1/2-GEMS and CDH-3-TF regulate SEAP and Luciferase expression, respectively. Drug concentrations ranged from 0.001 nM to 1 μM and different combinations were added depending on the protein components, Drug-1 + Drug-3 (b) and Drug-2 + Drug-3 (c). **d)** Scheme of multi-inputs single-output (MISO) designer cells where two drugs control one output. **e, f)** Quantification of SEAP activity controlled by CDH-1-TF and CDH-3-TF circuits (e) or CDH-1-GEMS and CDH-3-GEMS (f). Drug-3 concentrations ranged from 0.001 nM to 1 μM and Drug-1 treatments were performed in the presence of 1 μM Drug-3. **g)** Scheme of single-input multi-outputs (SIMO) designer cells where one drug controls two outputs. **h,i)** Quantification of SEAP and Luciferase activities under Drug-1 (h) or Drug-2 (i). AIR-1/2-GEMS coupled with SEAP expression circuits were co-transfected with CDH-1/2-TF circuits which control the Luciferase production. Respective drug concentrations ranged from 0.001 nM to 1 μM. All values presented are mean ± s.d. of three replicates and curves were fitted by four-parameters nonlinear regression.

We then engineered MISO cells (Fig. 5d) with CDH-1-TF and CDH-3-TF dual switches (Fig. 5e) that encode OFF behaviors controlled by Drug-1 and Drug-3. As expected, the individual CDHs showed similar response curves to those observed when they were tested in isolation (Fig. 2c). Interestingly, when treated with both drugs the engineered cells showed a unique behavior where the MISO’s response curve showed three persistent response levels (high, medium, low) controlled by drug amounts (Fig. 5e). This output behavior is distinct from the typical SISO systems which only show two persistent states (high, low) (Fig. 2c, f). Similarly, the engineered cells endowed with CDH-1-GEMS and CDH-3-GEMS dual-circuits (Fig. 5f) showed three persistent response levels. This type of output in biological systems could be utilized to produce graded levels of responses that are robust to minor oscillations in drug concentrations, and could be used to dissect fundamental biological processes but also to provide persistent response levels in translational applications.

Finally, we engineered SIMO cells (Fig. 5g) using two designed switches that produce ON and OFF responses upon exposure to the same drug. We transfected cells with combinations of AIR-GEMS and CDH-TFs which controlled the expression of the reporter proteins SEAP and Luciferase, respectively. Both Drug-1 and Drug-2 simultaneously activated SEAP production and suppressed the expression of Luciferase (Fig. 5h, i) in their respective switch systems, demonstrating that we successfully designed protein components which can physically dissociate and associate under the control of the same drug. To the best of our knowledge, Drug-1 and Drug-2 are the only two small molecules that like rapamycin can control both assembly^4,45^ and disassembly^15^ of protein components. Therefore, computational design expanded the repertoire of protein switches controlled by the same compounds that can be used to simultaneous turn ON and OFF synergistic outputs.

In summary, leveraging our computationally designed switches we created cellular bio-computing units with multiple output modalities that can be useful to deal with the inherent complexity of engineering mammalian cells and have the potential for translational applications in synthetic biology.

## Discussion

Protein components to chemically control cellular activities are the basis of many applications in synthetic biology^1^. However, the panel of protein switches available to control the assembly of protein complexes for synthetic biology and cell-based therapy applications remains extremely limited. Here, we presented a structure-based design blueprint to repurpose PPIs with known inhibitors into protein switches that can mediate both assembly and disassembly of hetero-dimeric complexes.

Specifically, we presented three CDH switches, controlled by three different drugs, along with multiple tailor-made affinity variants. Some of our observations on tuning CDH affinities highlight the challenge of designing components for acting in complex biological systems. For instance, by weakening the CDH interface, we can tune the affinity of the dimer down to 5,000-fold weaker. Yet despite this substantial difference in binding affinity, we find that the changes in cell-based measurements are much smaller (11 to 26-fold). Thus, we find a non-linear relationship between hetero-dimer affinities and drug IC_50_s in cells, pointing to the need to test a range of switches to optimize drug responsiveness. The CDHs can be utilized in the context of diverse proximity-based signal components intra- and extra-cellularly, which shows their modularity and wide applicability. Furthermore, CDH-2 (Bcl2:LD3) is controlled by an FDA-approved drug, Venetoclax and was optimized toward lower drug demand, which could reduce risks of drug-induced toxicity or off-target effects. This repurposed utilization of pre-clinical or FDA-approved drugs could open up the use of protein switches in the context of clinically relevant therapies.

In an effort to create chemically mediated dimerization switches, we further devised a multidomain architecture (referred to as AIR switches), for which we computationally designed a drug insensitive receptor that retained the ability to bind with the designed protein binder. The AIR switches showed an exquisite response to the drug inputs and we foresee that this design strategy will be broadly applicable to design other chemically controlled heterodimers, enriching this important class of protein components in synthetic biology. This expanded repertoire of CDHs and AIRs provides the opportunity to engineer more elaborated cellular outputs. As a proof-of-concept, our designed switches were combined in engineered cells which displayed several multi-input/multi-output behaviors. One interesting control mode was achieved where a single input drug regulated multiple outputs by exploiting the same smallmolecule to achieve ON- and OFF-responses simultaneously. This type of protein circuitry where the same small molecule can toggle between two antagonistic cellular outputs, may be a promising option for instance to control safety and efficacy in therapeutically relevant contexts.

Altogether, CDHs and AIRs enable a generalizable approach to develop controllable modules with the required properties for synthetic biology applications. We envision that the computational protein design toolbox will help to bridge the gap between the engineering of protein components and next-generation cell-based therapies.

## Contributions

SS, PG and BEC conceived the work. SS, PG, LS and BEC designed the experiments and analysed the results. PG performed computational design of CDHs and developed the protocol for multistate design. PG and SS performed the computational predication of AIRs. SS performed the experimental characterization. CY and YK solved x-ray structures. SR and SG supported the experimental work. SS, PG, LS and BEC wrote the manuscript, with input from all authors.

## Data availability

The data supporting the findings of this study are available within the article and its Supplementary Information. Coordinates of the determined structure have been deposited in the PDB with accession code 7AYE. All code used to perform the design simulation can be found at: https://github.com/LPDI-EPFL/CDH_AIR. Plasmids encoding the CDH and AIR components available from Addgene. Other data and reagents are available from the corresponding authors upon reasonable request.

## Acknowledgements

We thank Wanze Chen from the Deplancke laboratory for critical discussions. We thank Dominik Niopek, Fabian Sesterhenn and Raphael Brisset Di Roberto for comments in the manuscript. We thank Kelvin Lau and Florence Pojer from the PTPSP facility at EPFL for experimental support in protein expression and characterization. The computational simulations were facilitated by SCITAS at EPFL and by the Swiss National Supercomputer Center.

