## Supplementary material for "A rational blueprint for the design of chemically-controlled protein switches": supp material

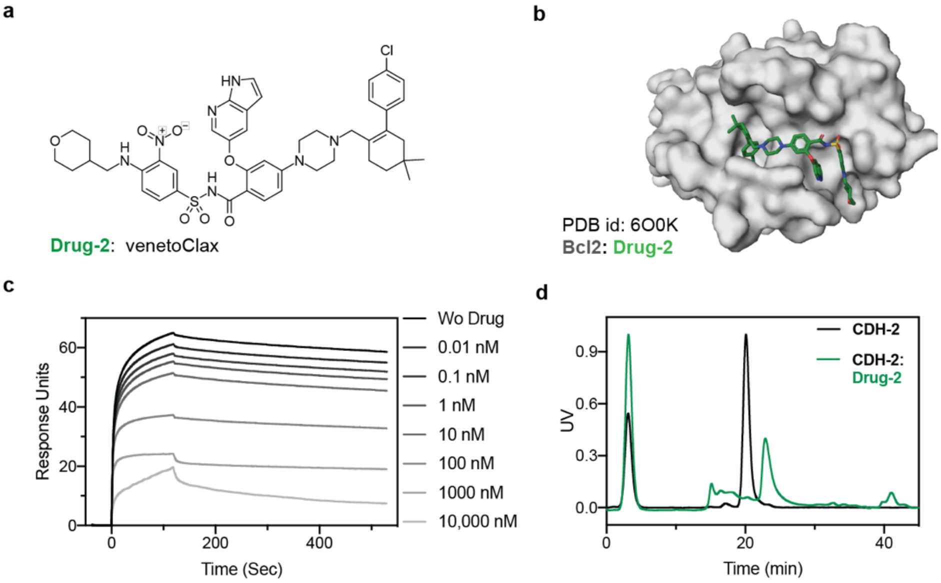

**Supplementary Figure 1: Biophysical characterization of CDH-2.**

**a)** Chemical structure of Venetoclax, used as CDH-2 disruptor (Drug-2). **b)** Crystal structure of Bcl2 (white surface) bound to Drug-2 (green sticks). **c)** Surface plasmon resonance drug competition assay. Kinetic curves of Drug-2 dissociating Bcl2:LD3 complex in SPR. Drug concentrations from 0.01 nM to 10 μM were pre-incubated with 4 μM LD3, then the mixtures were injected over a Bcl2-immobilized chip. Response units reflect the remaining interaction of Bcl2 and LD3 in the presence of serial diluted Drug-2. **d)** SEC-MALS analysis of CDH-2 showing Bcl2:LD3 complex with DMSO (black trace) and Bcl2:LD3 with Drug-2 did not result in complex formation and monomeric proteins (green trace), Bcl2 (19 kDa) and LD3 (16 kDa) eluted around 22 minutes.

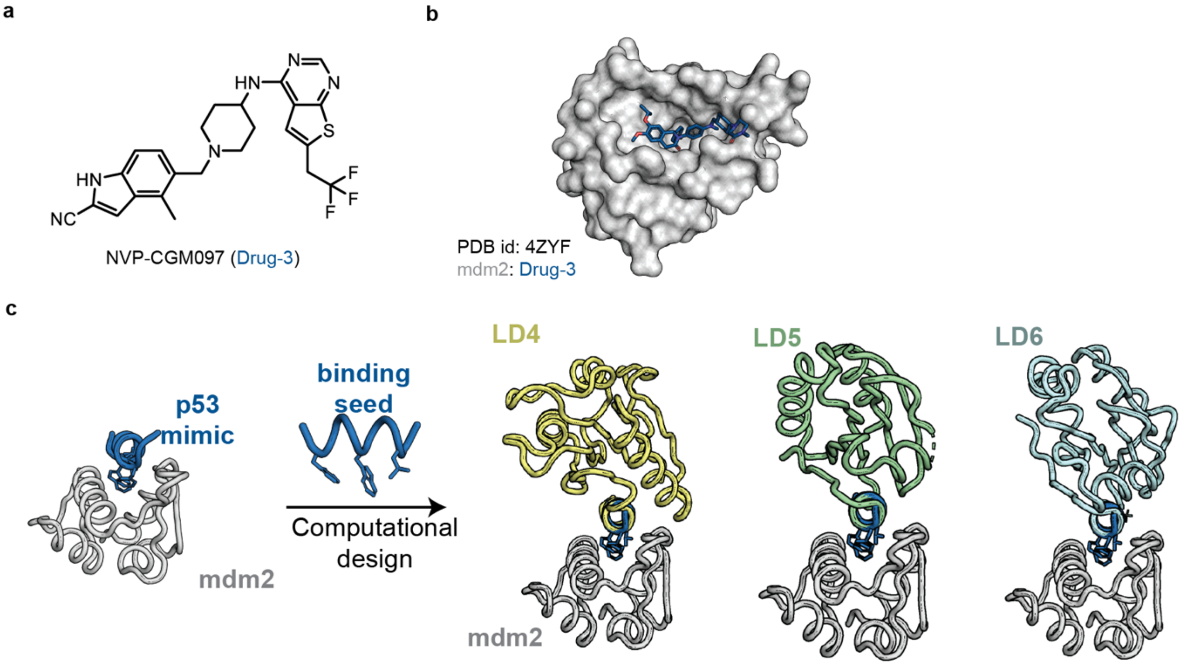

d

| Design id | LD4 | LD5 | **LD6** |
|---|---|---|---|
| Scaffold PDB id | 1AKU | 2IFQ | 2FWE |
| Scaffold protein name | Flavodoxin | Flavodoxin | DsbD |
| Organism of scaffold | *D. vulgaris* | *H. sapiens* | *M. musculus* |
| Rosetta ddG | -7.6 | -8 | -10 |
| Globularity | 1.04 | 0.98 | 0.93 |
| VdW Dots to scaffold | 980 | 660 | 670 |
| SASA of seed | 1769 | 1342 | 1337 |
| Total # Mutations on scaffold | 10 | 4 | 7 |

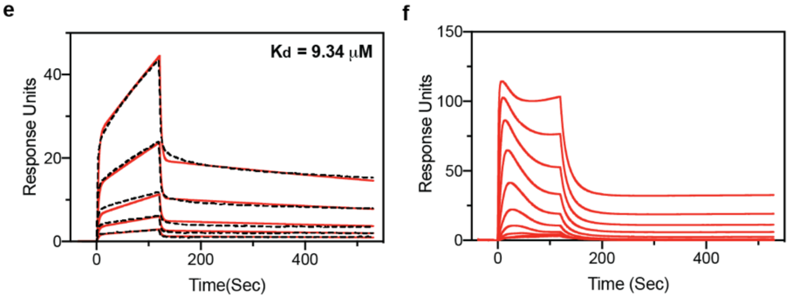

**Supplementary Figure 2: Computational design and experimental testing of mdm2 binders.**

**a)** Crystal structure of mdm2 in complex with its inhibitor:NVP-CGM097 (Drug-3). **b)** Chemical structure of Drug-3. **c)** p53 peptide was selected from the p53-mdm2 complex and shortened into an 8-residue motif which was matched against a database of > 11000 proteins using the MotifGraft protocol. Structures of three candidate designs (LD4-6) are shown in complex with mdm2. **d)** Table of designs and scores for the scoring/filtering criteria. Scaffold PDB ID: Protein Data Bank ID of the protein that was used as a scaffold to design each binder. Scaffold protein name: Name of the protein used as a scaffold. Organism of scaffold: Species origin of the native protein. Rosetta ddG: Computed delta-delta G interaction energy between designs and mdm2. Globularity: Globularity score for each designed scaffold, where values closer to 1.0 are more globular. Globularity score was based on a metric created by Miller et al[1], further explained in Methods. vdW Dots to scaffold: Number of Van der Waals (VdW) contacts between the grafted motif and the scaffold. SASA of seed: buried surface area of the grafted motif in the scaffold. Total # mutations on scaffold: final number of residues in the scaffold that were mutated during the design process. **e-f)** Affinity measurements of mdm2 and designed binders: LD4 (e), LD5 (f). Designed binders at concentrations from 125 to 2000 nM with 2-fold dilutions were injected over mdm2 immobilized chips. Black dashed curves represent the sensorgrams and the fitting curves are in solid red curves. $K_d$s were computed using a 1:1 binding model.

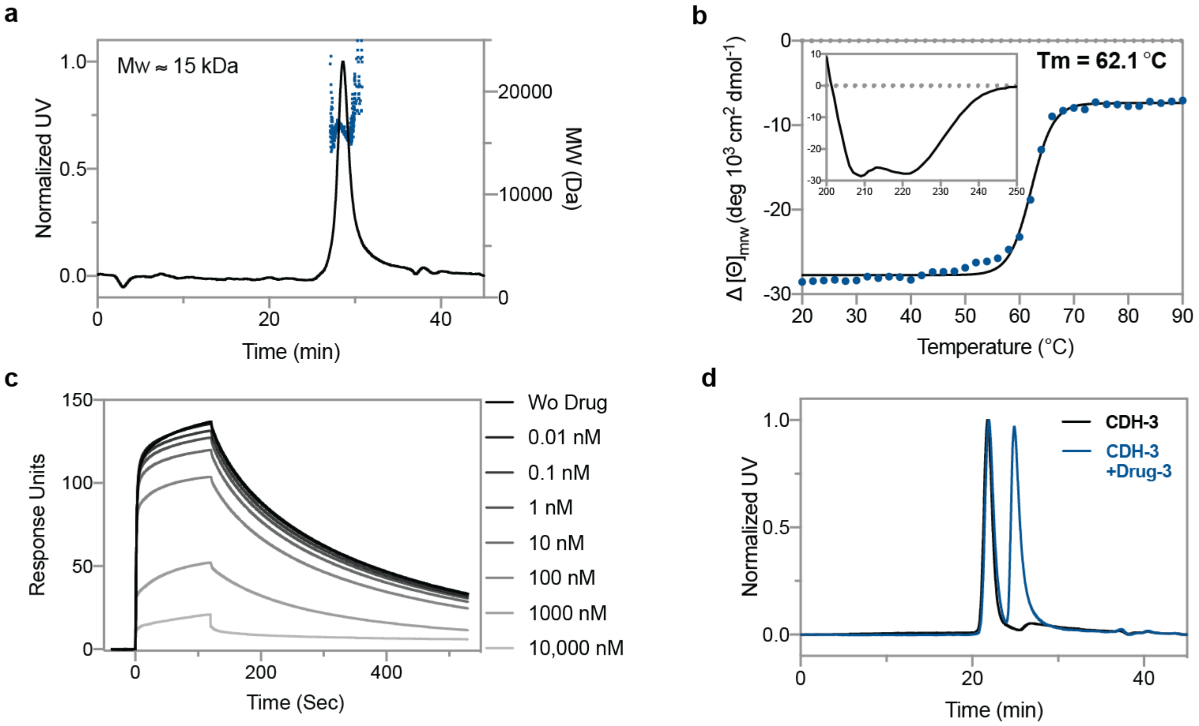

**Supplementary Figure 3: Biophysical characterization of CDH-3.**

**a)** SEC-MALS analysis showed that LD6 is a monomeric protein with a molecular weight around 15 kDa. **b)** Thermostability of LD6. Circular Dichroism spectrum showed a melting temperature of 62 ˚C and a typical helical absorption curve at 220 nm. **c)** Surface plasmon resonance drug competition assay. Kinetic curves of Drug-3 dissociating mdm2:LD6 complex in SPR. Drug concentrations from 0.01 nM to 10 μM were pre-incubated with 4 μM LD6, then the mixtures were injected over an mdm2-immobilized chip. Response units reflect the remaining interaction of mdm2 and LD6 in the presence of serially diluted Drug-3. **d)** SEC-MALS analysis of CDH-3 showing mdm2:LD6 complex with DMSO (black trace), while dissociation of the mdm2:LD6 with Drug-3 could be observed and monomeric proteins (blue trace), mdm2 (11 kDa) and LD6 (15 kDa) eluted around 25 minutes.

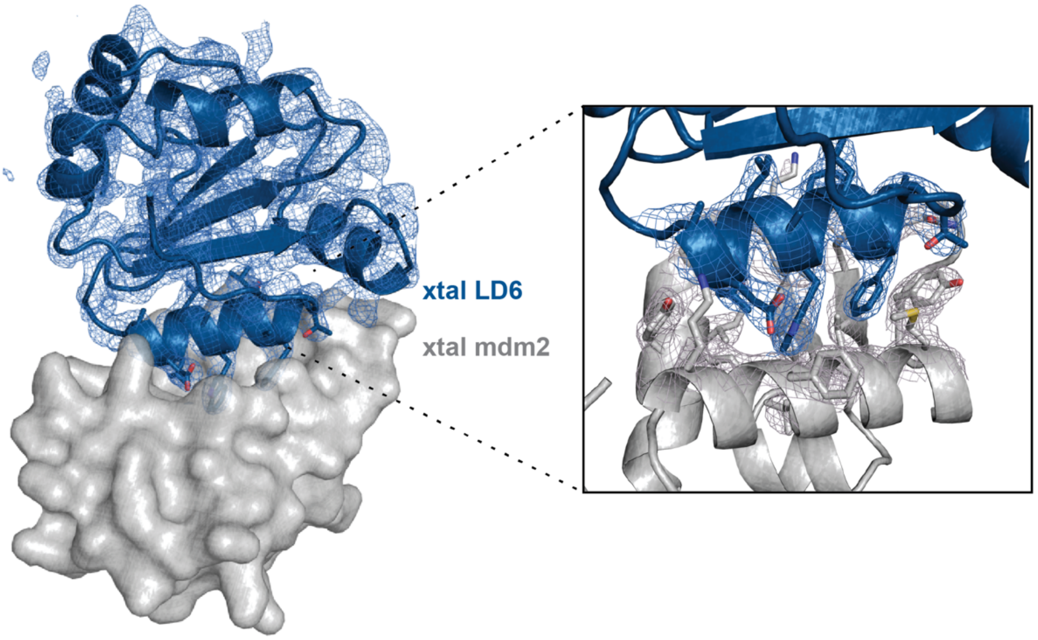

**Supplementary Figure 4: 2mFo-mFc electron density maps of the mdm2:LD6 crystal structure.** Maps are contoured at 1 σ and carved around the structure at 1.6 Å. Side chains are shown in sticks representation.

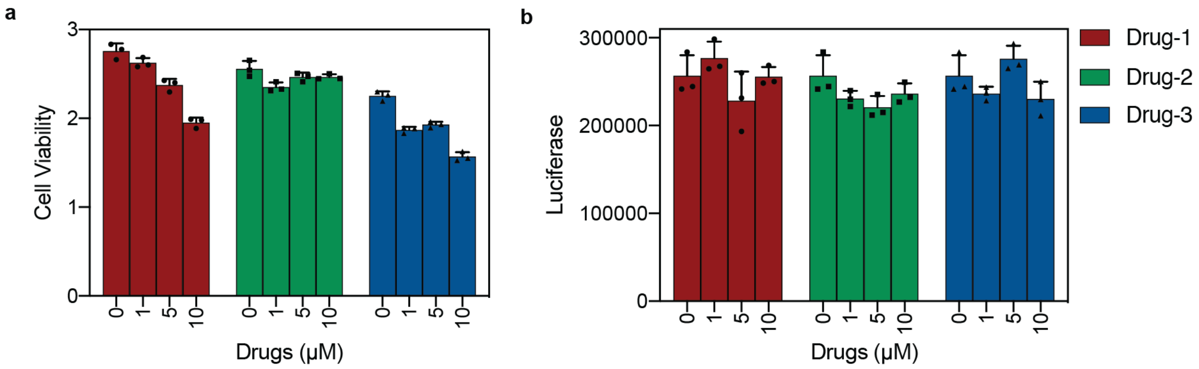

**Supplementary Figure 5: Drug toxicity assessment.**

**a)** Effect of Drugs-1-3 on the viability of HEK293T cells. Absorbance of CCK8 measurement at 450 nm with indicated drug concentrations. Each bar represents the mean of three biological replicates ± s.d.
**b)** Effect of drug on HEK293T cells. Quantification of Luciferase activity 24 hours after the addition of the drugs. The HEK293T cells were transfected with the Luciferase reporter and constitutive Gal4-Rel65 expression plasmid (pCMV-Gal4-Rel65-pA).

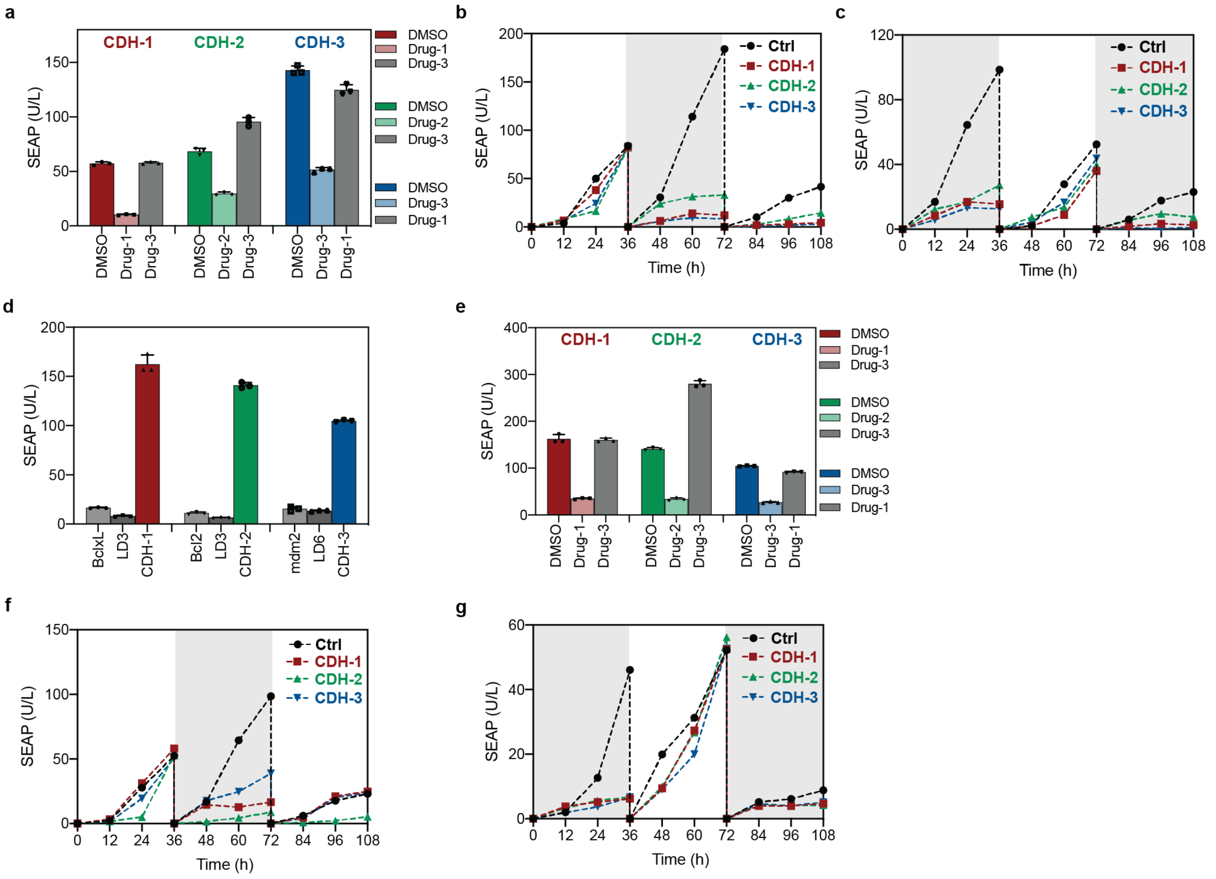

**Supplementary Figure 6: Dose-dependence and reversibility characterization of CDHs in CDH-TF and CDH-GEMS systems.**

**a)** Drug specificity of CDH-TFs. Quantification of SEAP activity of three CDHs with DMSO (negative control; dark color), specific (designed drug induced OFF switches; light color) and unspecific drug treatment (testing for possible unintended drug induced effects; gray). **b)** Dynamic regulation of SEAP expression mediated by CDHs-TF in modes of ON-OFF-ON. The control samples (Ctrl) were transfected with the SEAP reporter S132 and constitutive Gal4-Rel65 expression plasmid S108 (pCMV-Gal4-Rel65-pA). During the OFF-time period shown in gray box, CDH-TF cells were cultured with 500 nM of the corresponding drugs while Ctrl cells were treated with same concentration of DMSO, drugs/DMSO were replenished every 12 hours at time points of 36h, 48h and 60h. Cells were split at time points of 36h and 72h which changed medium to with/without drugs respectively. **c)** Dynamic regulation of SEAP expression mediated by CDHs-TF in modes of OFF-ON-OFF. Time point 0 was set 12 hours post transfection to start the intermittent drug treatment shown in gray boxes, CDH-TF cells were cultured with 500 nM of the corresponding drugs while Ctrl cells were treated with same concentration of DMSO. Drugs/DMSO were refreshed at time points of 0h, 12h, 24h and 72h, 84h, 96h. Cells were split at time points of 36h and 72h which changed medium to without/with drugs respectively. **d)** SEAP (secreted embryonic alkaline phosphatase) activity upon transfecting single or paired constructs of CDH-GEMS. Single constructs in gray remain inactive, while paired transfection of CDHs (1-red, 2-green and 3-blue) activate SEAP expression. **e)** Drug specificity test on CDH-GEMS.

Quantification of SEAP activity of three CDHs with DMSO (dark color), specific (light color) and unspecific drug treatment (gray). Samples of CDHs (1-red, 2-green and 3-blue) were treated with their specific disruptors in faded color labelled as Drug-1, Drug-2 and Drug-3, and the unspecific Drug-3 used for CDH-1 and CDH-2, Drug-1 as the unspecific disruptor for CDH-3. **f-g)** Dynamic regulation of SEAP expression mediated by CDH-GEMS. Positive control was transfected with CDH-1-GEMS and with DMSO treatment. Experiments were done with the same principle in panel b and c. Samples (culture medium) in panels of b, c, f and g were collected every 12 hours. Values were normalized to zero based on the SEAP expression at time 0.

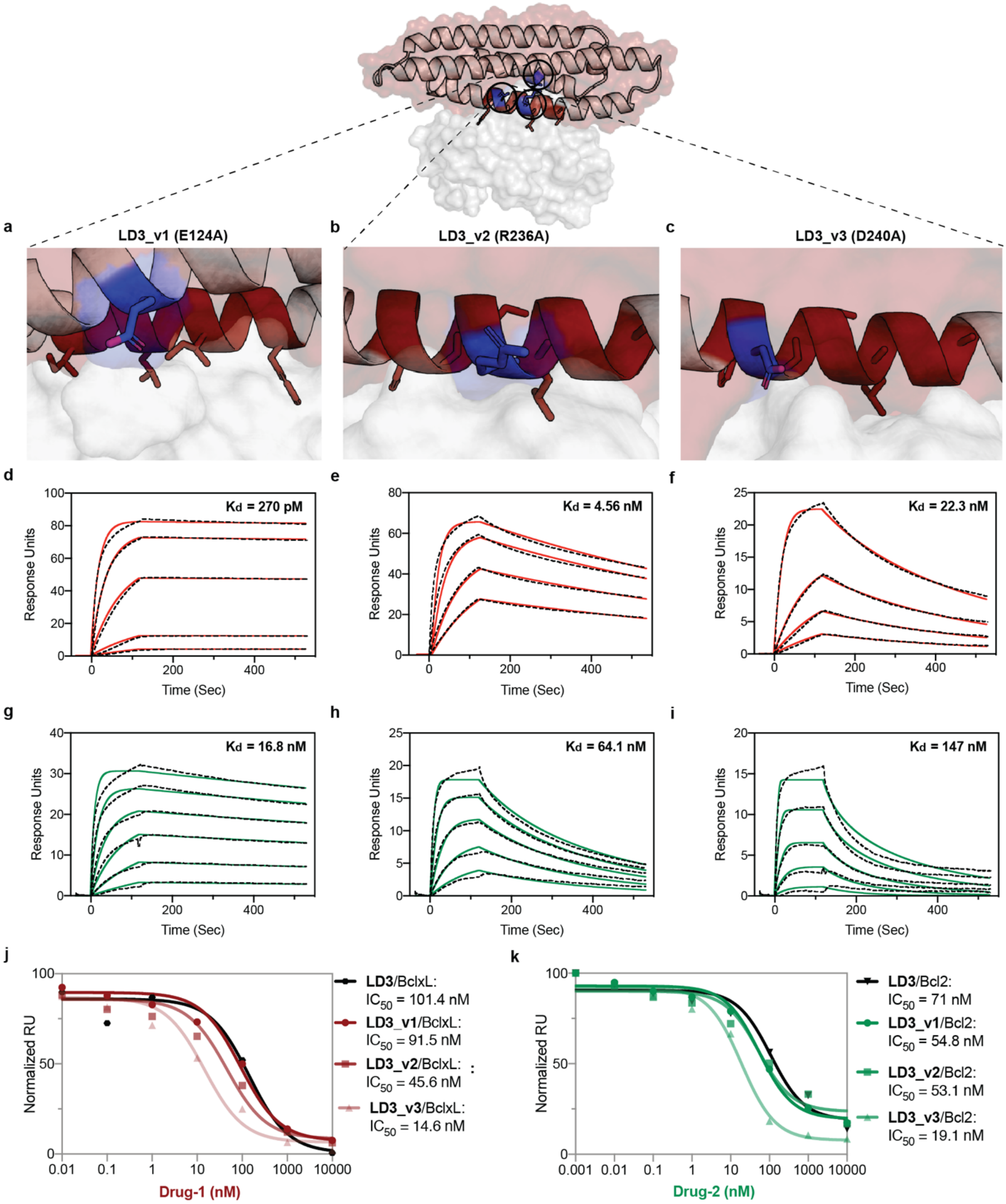

**Supplementary Figure7: Evaluation of weak affinity variants of the LD3:BclxL and LD3:Bcl2 complexes.**

**a-c)** Alanine mutations of LD3_v1, LD3_v2 and LD3_v3. The LD3 protein (light red cartoon and surface) is shown in complex with Bcl-XL protein (white surface). A 12-amino acid motif from the BiM BH3 peptide (dark red cartoon) is highlighted with hotspot residues shown in red sticks and with each residue mutated to alanine shown in blue sticks. **d-f)** Affinity measurements of Bcl-XL and LD3 (v1_v3) by SPR. Indicated concentrations of LD3 mutants were injected over the Bcl-XL immobilized chips. The binding sensorgrams (black dashed curves) are plotted with fitted curves (solid red). **g-i)** Affinity measurements of Bcl2 and LD3 (v1_v3) by SPR. Indicated concentrations of LD3 mutants were injected over the Bcl2 immobilized chips. The binding sensorgrams (black dashed curves) are plotted with fitted curves (solid green). **d-i)** $K_d$ was calculated by a 1:1 binding model. **j-k)** SPR $IC_{50}$ determinations of LD3 mutants with Bcl-XL (**j**) and Bcl2 (**k**). Indicated concentrations of Drug-1 or Drug-2 were premixed with 4 μM LD3 and mutants, then injected into Bcl-XL and Bcl2 immobilized chips, respectively. Readouts from the sensorgrams were extracted at the time point 120 s to calculate the $IC_{50}$s using a three-parameter nonlinear regression.

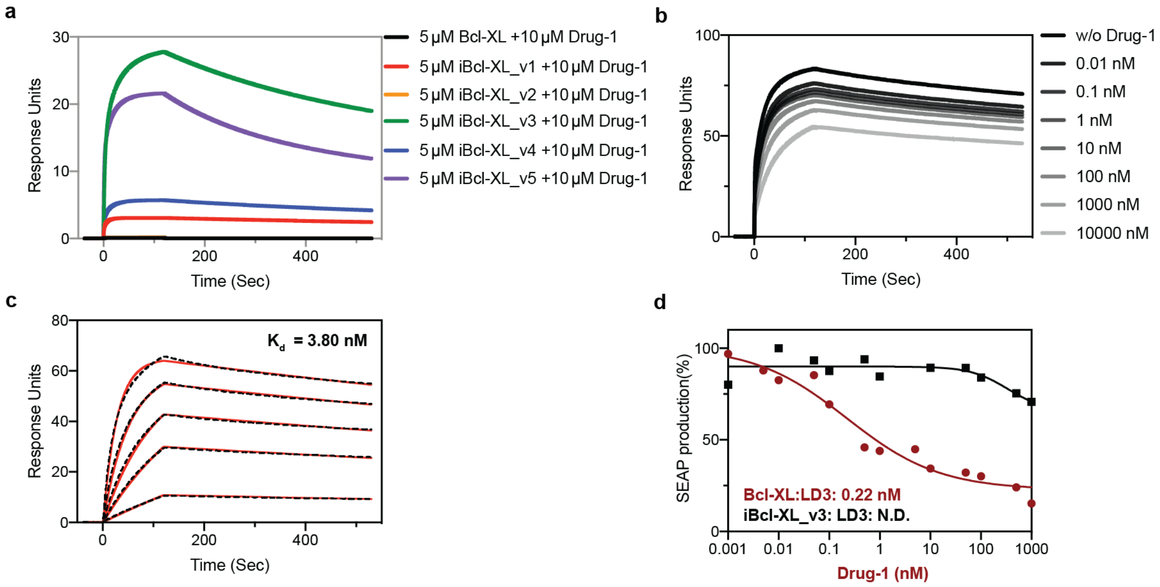

**Supplementary Figure 8: Biophysical and cellular characterization of designed Drug-1-resistant BclxL mutants.**

**a)** Pre-screening of Drug-1 resistant Bcl-XL mutants by SPR. 5 μM of Bcl-XL and five mutants were mixed with 10 μM of Drug-1 and injected over the LD3 immobilized chip to analyse the binding response. Drug-1 resistant mutants showed higher response. Drug-1 showed complete inhibition to Bcl-XL (Black) and iBcl-XL_v2 (orange; not visible in the graph due to overlay with Bcl-XL), significant inhibition to iBcl-XL_v1 (red) and iBcl-XL_v4 (blue), mild inhibition to iBcl-XL_v5 (purple) and the weakest inhibition to iBcl-XL_v3 (green). **b)** SPR competition assay of Drug-1 dissociating iBcl-XL_v3:LD3 complex. Serial dilutions of Drug-1 were pre-mixed with 4 μM LD3 and injected over iBcl-XL_v3 immobilized chip to collect the response. **c)** Dissociation constant measurement of iBcl-XL_v3 and LD3. Sensorgrams are in black dashed curves and the fitted curves in solid red lines. $K_d$ was using a 1:1 binding model. **d)** $IC_{50}$s of Drug-1 dissociating Bcl-XL:LD3 versus iBcl-XL_v3:LD3 in the CDH-GEMS system. The plot shows the dose response of 24 hours after the addition of serial Drug-1 concentrations. Values were normalized to the positive control (DMSO group). Each data point represents the mean of n = 3 biological replicates, and the $IC_{50}$s were calculated using four-parameter nonlinear regression.

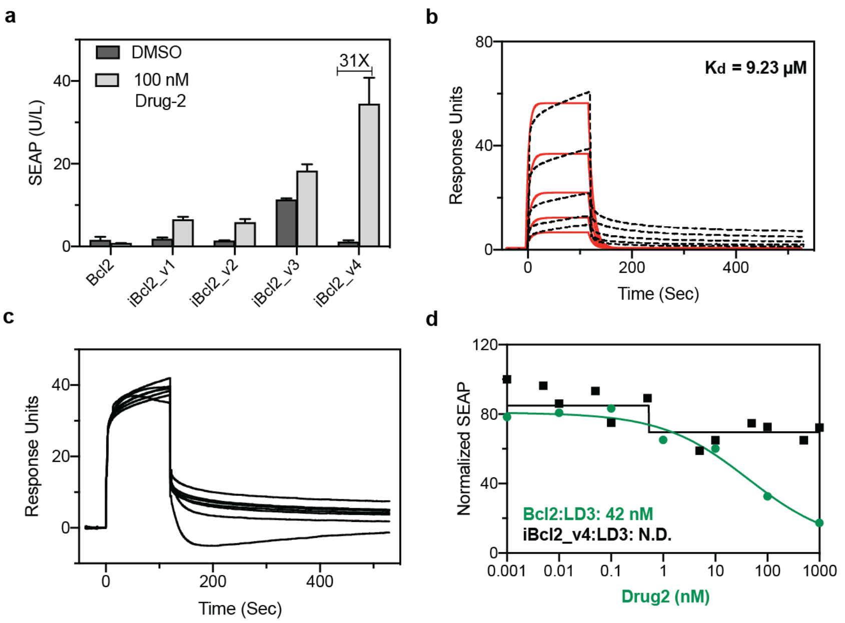

**Supplementary Figure9: Cellular and biophysical characterization of Drug-2 resistant Bcl2 mutants.**

**a)** Pre-screening of Drug-2 resistant Bcl2 mutants. Bcl2 mutants were cloned into the AIR-GEMS platform and co-transfected with $P_{SV40}$-IgK-Bcl2-$GGGGS_{X3}$-LD3-EpoRm-IL-6RBm-pA to test ON-switch behavior. $P_{SV40}$-IgK-Bcl2-EpoRm-IL-6RBm-pA was used as the negative control, and all groups were exposed to 100nM DMSO (dark grey) versus Drug-2 (light grey). **b)** Dissociation constant measurement of iBcl2_v4 and LD3 in SPR. Sensorgrams are in black dashed curves and the fitted curves in solid red lines. $K_D$ was calculated by a 1:1 binding model in the biacore system. **c)** Drug competition assay measured Drug-2 disrupting iBcl2_v4:LD3 complex. Serial dilutions of Drug-2 were mixed with 4 μM LD3 and injected over iBcl2_v4 immobilized chip to collect the response. **d)** $IC_{50}$s of Drug-2 disrupting Bcl2:LD3 and iBcl2_v4:LD3 in the CDH-GEMS system. Dose response of diluted Drug-2, 24 hours after treatment. Each data point represents the mean of n = 3 biological replicates, and the $IC_{50}$s were calculated using four-parameter nonlinear regression.

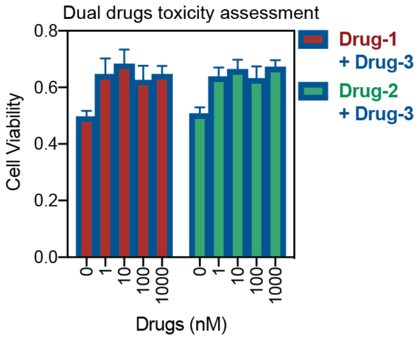

**Supplementary Figure 10: Dual-drug toxicity test on 293T cells.**

Dual drug toxicity tests in 293T cells. Drug-1 + Drug-3 and Drug-2 + Drug-3 were diluted to the concentrations of 1 nM, 10 nM, 100 nM and 1000 nM each drug in cell culture medium. CCK8 assay was performed 24 hours after drug treatment.

**Supplementary Table 1: Crystal structure information of mdm2: LD6**

| | **LD6/mdm2** |
|---|---|
| **Data collection** | |
| Space group | P $4_3$ 2 2 |
| Cell dimensions | |
| *a*, *b*, *c* (Å) | 73.2, 73.2, 92.2 |
| α, β, γ (°) | 90.0, 90.0, 90.0 |
| Resolution (Å) | 45.16 – 2.95 (3.13– 2.95)* |
| *I* / σ*I* | 22.2 (2.2) |
| Completeness (%) | 99.8 (99.8) |
| Redundancy | 6.9 (7.4) |
| $R_{meas}$ | 0.07 (0.84) |
| CC1/2** | 0.99 (0.67) |
| **Refinement** | |
| No. reflections | 5,646 |
| $R_{work}$ / $R_{free}$ | 0.20 / 0.25 |
| No. atoms | 1,656 |
| Protein | 1,656 |
| *B*-factors | |
| Protein | 86.1 |
| R.m.s. deviations | |
| Bond lengths (Å) | 0.008 |
| Bond angles (°) | 1.61 |
| **PDB code** | **7AYE** |

*Values in parentheses are for highest-resolution shell.

**CC1/2 refers to Pearson's correlation coefficients (CC) between intensity estimates from half data sets.

**Supplementary Table 2: Prediction of Drug-1 resistant Bcl-XL mutants.**

**Mutations**: Specific mutations modeled from wildtype Bcl-XL. Positive_Bound: Score of the LD3:iBcl-XL mutant complex.

**Positive_unbound**: Score of the iBcl-XL mutant in its unbound (apo) state.

**Negative_Bound**: score of the Drug-1:iBcl-XL complex.

**Negative_unbound:** Score of iBcl-XL mutant in its unbound (apo) state, based on the crystal structure of Drug-1:Bcl-XL.

**Ratio_unbound**: (Positive_bound - Positive_unbound) - (Negative_bound - Negative_unbound).

**Ratio_Bound:** (Positive_bound - Negative_bound)

| Name | Mutations | Positive_Bound | Positive_unbound | Negative_Bound | Negative_unbound | Ratio_unbound | Ratio_Bound |
|---|---|---|---|---|---|---|---|
| iBcl-XL_v1 | T109L, A149L | -362.525 | -164.568 | -130.467 | -155.829 | -223.319 | -232.058 |
| iBcl-XL_v2 | A149V | -366.699 | -139.013 | -180.803 | -150.961 | -197.597 | -185.649 |
| iBcl-XL_v3 | R102E, F105I | -365.076 | -161.623 | -170.317 | -149.164 | -182.3 | -194.759 |
| iBcl-XL_v4 | R102F, T109V | -368.015 | -164.292 | -174.012 | -146.991 | -176.702 | -194.003 |
| iBcl-XL_v5 | E98S, F105I | -371.963 | -160.491 | -172.324 | -148.471 | -187.619 | -199.639 |

**Supplementary Table 3: Prediction of Drug-2 resistant Bcl2 mutants.**

**Mutations**: Specific mutations modeled from wildtype Bcl2. Positive_Bound: Score of the LD3:iBcl2 mutant complex.

**Positive_unbound**: Score of the iBcl2 mutant in its unbound (apo) state.

**Negative_Bound**: score of the Drug-2:iBcl2 complex.

**Negative_unbound:** Score of iBcl2 mutant in its unbound (apo) state, based on the crystal structure of Drug-2:Bcl2.

**Ratio_unbound**: (Positive_bound - Positive_unbound) - (Negative_bound - Negative_unbound).

**Ratio_Bound:** (Positive_bound - Negative_bound)

Three additional mutations were enriched in the top designs, and therefore we designed a further version, iBcl2_v4 (100V_103N_202H).

| Name | Mutations | Positive_Bound | Positive_unbound | Negative_Bound | Negative_unbound | Ratio_unbound | Ratio_Bound |
|---|---|---|---|---|---|---|---|
| iBcl2_v1 | V156I_Y202H | -383.197 | -207.19 | -223.495 | -207.19 | -159.702 | -159.702 |
| iBcl2_v2 | D103N_Y202H | -373.831 | -193.74 | -222.264 | -193.74 | -151.567 | -151.567 |
| iBcl2_v3 | A100T_D103S | -371.663 | -191.485 | -220.949 | -191.485 | -150.714 | -150.714 |
| iBcl2_v4 | 100V_103N_202H | | | | | | |

**Supplementary Table 4: Protein sequences of CDHs(1-3).**

| Name | Sequences |
|---|---|
| CDH-1-Bcl-XL | MSQSNRELVVDFLSYKLSQKGYSWSQFSDVEENRTEAPEGTESEAVKQALREAGDEFELRYRRAFSDLTSQLHITPGTAYQSFEQVVNELFRDGVNWGRIVAFFSFGGALCVESVDKEMQVLVSRIAAWMATYLNDHLEPWIQENGGWDTFVELYGNNAAAESRKGQER |
| CDH-2-Bcl2 | MAHPGRTGYDNREIVMKYIHYKLSQRGYEWDAGDDVEENRTEAPEGTESEVVHLTLRQAGDDFSRRYRRDFAEMSSQLHLTPFTARGRFATVVEELFRDGVNWGRIVAFFEFGGVMCVESVNREMSPLVDNIALWMTEYLNRHLHTWIQDNGGWDAFVELYGPSMR |
| CDH-1/2-LD3 | QRWELALGRFLEYLSWVSTLSEQVQEELLSSQVTQELRALMDETMKELKAYKSELEEQLTPVAEETRARLSKELQAAQARLGADMEDVRGRLVQYRGEVQAMLGQSTEELRVRLASHLIALALRLIGDAFDLQKRLAVY |
| CDH-3-mdm2 | GPLGSSQIPASEQETLVRPKPLLLKLLKSVGAQKDTYTMKEVLFYLGQYIMTKRLYDAAQQHIVYCSNDLLGDLFGVPSFSVKEHRKIYTMIYRNLV |
| CDH-3-LD6 | HLNFTQIKTAFALYWALLEAQGKPVMLDLYADWCVACKEFEKYTFSDPQVQKALADTVLLQANVTANDAQDVALLKHLNVLGLPTILFFDGQGQEHPQARVTGFMDAETFSAHLRDRQPHHH |

**Supplementary Table 5: LD3 variants with low affinity and drug-insensitive receptors.**

| Name | Sequences |
|---|---|
| LD3_v1 | QRWELALGRFL**A**YLSWVSTLSEQVQEELLSSQVTQELRALMDETMKELKAYKSELEEQLTPVAEETRARLSKELQAAQARLGADMEDVRGRLVQY<br>RGEVQAMLGQSTEELRVRLASHLIALALRLIGDAFDLQKRLAVY |
| LD3_v2 | QRWELALGRFLEYLSWVSTLSEQVQEELLSSQVTQELRALMDETMKELKAYKSELEEQLTPVAEETRARLSKELQAAQARLGADMEDVRGRLVQY<br>RGEVQAMLGQSTEELRVRLASHLIALAL**A**LIGDAFDLQKRLAVY |
| LD3_v3 | QRWELALGRFLEYLSWVSTLSEQVQEELLSSQVTQELRALMDETMKELKAYKSELEEQLTPVAEETRARLSKELQAAQARLGADMEDVRGRLVQY<br>RGEVQAMLGQSTEELRVRLASHLIALALRLIG**A**AFDLQKRLAVY |
| iBcl-XL_v3 | MSQSNRELVVDFLSYKLSQKGYSWSQFSDVEENRTEAPEGTESEAVKQALREAGDEFELRY**E**RA**I**SDLTSQLHITPGTAYQSFEQVVNELFRDGV<br>NWGRIVAFFSFGGALCVESVDKEMQVLVSRIAAWMATYLNDHLEPWIQENGGWDTFVELYGNNAAAESRKGQER |
| iBcl2_v4 | MAHPGRTGYDNREIVMKYIHYKLSQRGYEWDAGDDVEENRTEAPEGTESEVVHLTLRQ**V**GD**N**FSRRYRRDFAEMSSQLHLTPFTARGRFATVVEE<br>LFRDGVNWGRIVAFFEFGGVMCVESVNREMSPLVDNIALWMTEYLNRHLHTWIQDNGGWDAFVEL**H**GPSMR |

**Supplementary Table 6: Plasmids in cellular applications**

| Plasmid | Description and cloning strategy | Reference |
|---|---|---|
| pPKm-118 | $P_{UAS}$-driven vector expressing reporter gene Luciferase ($P_{5XUAS}$-Luciferase-pA). | Addgene #90491 |
| S132 | $P_{UAS}$-driven vector expressing reporter gene secreted embryonic alkaline phosphatase ($P_{5XUAS}$-SEAP-pA). | This work |
| S108 | Constitutive $P_{hCMV}$-driven mammalian expression vector ($P_{hCMV}$-Gal4-Rel65-pA), gene cloned from plasmid: pCS2+ Gal4-GBP2-IRES-GBP7-p65 | Addgene #50020 |
| S111 | Constitutive $P_{hCMV}$-driven mammalian expression vector of CDH-1-TF cassette ($P_{hCMV}$-Gal4-Bcl-XL-P2A-LD3-Rel65-pA), cloned into the NheI digested S108 by Gibson assembly | This work |
| S112 | Constitutive $P_{hCMV}$-driven mammalian expression vector of CDH-2-TF cassette ($P_{hCMV}$-Gal4-Bcl2-P2A-LD3-Rel65-pA), cloned into the NheI digested S108 by Gibson assembly | This work |
| S113 | Constitutive $P_{hCMV}$-driven mammalian expression vector of CDH-3-TF cassette ($P_{hCMV}$-Gal4-mdm2-P2A-LD6-Rel65-pA), cloned into the NheI digested S108 by Gibson assembly | This work |
| pLS13 | Mammalian reporter plasmid for STAT3-induced SEAP expression (OStat3-$P_{hCMVmin}$-SEAP-pA) | This work |
| pLS15 | Constitutive $P_{hCMV}$-driven mammalian STAT3 expression vector ($P_{hCMV}$-STAT3-pA) | This work |
| S184 | Mammalian CDH-1-GEMS expression vector ($P_{SV40}$-IgK-Bcl-XL-EpoRm-IL-6RBm-pA), original plasmid was from pLeo619 by changing the extracellular interaction domain into Bcl-XL to form CDH-1 with LD3(S185), cloned into the SpeI digested pLEo619 by Gibson assembly with the secrection signal of IgK | This work |
| S193 | Mammalian CDH-2-GEMS expression vector ($P_{SV40}$-IgK-Bcl2-EpoRm-IL-6RBm-pA), co-transfection with S185 to form the full CDH-2-GEMS machinery | This work |
| S185 | Mammalian expression vector of binder protein LD3, used together with S184 and S193 to form CDH-1 and CDH-2 respectively ($P_{SV40}$-IgK-LD3-EpoRm-IL-6RBm-pA) | This work |
| S189 | Mammalian CDH-3-GEMS expression vector ($P_{SV40}$-IgK-mdm2-EpoRm-IL-6RBm-pA), work with S190 | This work |
| S190 | Mammalian CDH-3-GEMS expression vector ($P_{SV40}$-IgK-LD6-EpoRm-IL-6RBm-pA), work with S189 | This work |
| S215 | Mammalian AIR-1-GEMS expression vector ($P_{SV40}$-IgK-Bcl-XL-$GGGGS_{X3}$-LD3-EpoRm-IL-6RBm-pA), two domains of CDH-1 were genetically fused by the three repeats of GS linker and cloned into EpoR-IL-6RB receptor plasmid | [11] |

| Plasmid | Description and cloning strategy | Reference |
|---|---|---|
| S226 | Mammalian AIR-1-GEMS expression vector ($P_{SV40}$-IgK-iBcl-XL_v3-EpoRm-IL-6RBm-pA), the drug insensitive Bcl-XL variant, Bcl-XLMut3 was cloned into EpoR-IL-6RB receptor backbone and co-worked with S215 to form the AIR-1-GEMS switch | This work |
| S222 | Mammalian AIR-2-GEMS expression vector ($P_{SV40}$-IgK-Bcl2-$GGGGS_{X3}$-LD3-EpoRm-IL-6RBm-pA), two domains of CDH-1 were genetically fused by the three repeats of GS linker and cloned into EpoR-IL-6RB receptor plasmid | This work |
| S228 | Mammalian AIR-1-GEMS expression vector ($P_{SV40}$-IgK-iBcl2_v4-EpoRm-IL-6RBm-pA), the drug insensitive Bcl2 variant, Bcl2Mut7 was cloned into EpoR-IL-6RB receptor backbone and co-worked with S222 to form the AIR-2-GEMS switch | This work |

**Supplementary Table 7: transfection table of CDH and AIRs in cells.**

The amount of DNA was calculated per 1 well of a 96-well plate.

| Application | Reporter | Effector | Other plasmids | Total DNA |
|---|---|---|---|---|
| CDH-(1-3)-TF | S132<br>50 ng | S111/S112/S113<br>50 ng | None | 100 ng |
| CDH-(1-3)-GEMS | pLS13<br>30ng | S184 + S185<br>/S193+S185/S189+S190<br>50 ng + 50 ng | **pLS15**<br>**3.3ng** | 133.3 ng |
| AIR-(1-2)-GEMS | pLS13<br>30ng | S215 + S226<br>/S222 + S228<br>50 ng + 50 ng | **pLS15**<br>**3.3ng** | 133.3 ng |
| MIMO-Drug-1+Drug-3 | pPKm-118 30 ng<br>pLS13 50 ng | S113 30 ng<br>S215 + S226 50 ng + 50 ng | pLS15 3.3 ng | 193.3 ng |
| MIMO-Drug-2+Drug-3 | pPKm-118 30 ng<br>pLS13 50 ng | S113 30 ng<br>S215 + S226 50 ng + 50 ng | pLS15 3.3 ng | 193.3 ng |
| SIMO-Drug-1 | pPKm-118 30 ng<br>pLS13 50 ng | S111 30 ng<br>S215 + S226 50 ng + 50 ng | pLS15 3.3 ng | 193.3 ng |
| SIMO-Drug-2 | pPKm-118 30 ng<br>pLS13 50 ng | S112 30 ng<br>S215 + S226 50 ng + 50 ng | pLS15 3.3 ng | 193.3 ng |
| MISO-Drug-1 + Drug-3 | S132 30 ng<br>pLS13 50 ng | S113 30 ng<br>S215 + S226 50 ng + 50 ng | pLS15 3.3 ng | 193.3 ng |
| MISO-Drug-2 + Drug-3 | S132 30 ng<br>pLS13 50 ng | S113 30 ng<br>S215 + S226 50 ng + 50 ng | pLS15 3.3 ng | 193.3 ng |

**Supplementary Table 8: Summary of tested OFF/ON switches.**

| Name | Protein components | Disruptor/ Inducer | Corresponsing plasmids | Applications |
|---|---|---|---|---|
| OFF switches | | | | |
| CDH-1-TF | Bcl-XL & LD3 | Drug-1 | S111 | OFF switchable TF system |
| CDH-2-TF | Bcl2 & LD3 | Drug-2 | S112 | OFF switchable TF system |
| CDH-3-TF | mdm2 & LD6 | Drug-3 | S113 | OFF switchable TF system |
| CDH-1-GEMS | Bcl-XL & LD3 | Drug-1 | S184 & S185 | OFF switchable GEMS system |
| Weakened CDH-1-GEMS | Bcl-XL & LD3-v3 | Drug-1 | S184 & S188 | More senstitive OFF switchabe GEMS system |
| CDH-2-GEMS | Bcl2 & LD3 | Drug-2 | S193 & S185 | OFF switchable GEMS system |
| Weakened CDH-2-GEMS | Bcl2 & LD3-v3 | Drug-2 | S193 & S188 | More senstitive OFF switchabe GEMS system |
| CDH-3-GEMS | mdm2 & LD6 | Drug-3 | S189 & S190 | OFF switchable GEMS system |
| iCDH-1-GEMS | iBcl-XL_v3 & LD3 | Drug-1 | S226 & S185 | insensitive CDH-1 |
| iCDH-2-GEMS | iBcl2_v4 & LD3 | Drug-2 | S228 & S185 | insensitive CDH-2 |
| ON-switches | | | | |
| AIR-1-GEMS | iBcl-XL_v3 & CDH-1(BclxL-GS-LD3) | Drug-1 | S215 & S226 | ON switchable GEMS system |
| AIR-2-GEMS | iBcl2_v4 & CDH-2(Bcl2-GS-LD3) | Drug-2 | S222 & S228 | ON switchable GEMS system |

**Supplementary Table 9: Summary of control logics.**

| | | | |
|---|---|---|---|
| SISO | | | |
| CDH-1-TF | Drug-1 | SEAP/Luciferase | Fig. 3 |
| CDH-2-TF | Drug-2 | SEAP/Luciferase | Fig. 3 |
| CDH-3-TF | Drug-3 | SEAP/Luciferase | Fig. 3 |
| CDH-1-GEMS | Drug-1 | SEAP | Fig. 3 |
| Weakened CDH-1 | Drug-1 | SEAP | Fig. 3 |
| CDH-2-GEMS | Drug-2 | SEAP | Fig. 3 |
| Weakened CDH-2 | Drug-2 | SEAP | Fig. 3 |
| CDH-3-GEMS | Drug-3 | SEAP | Fig. 3 |
| iCDH-1-GEMS | | SEAP | Supp Fig. 8 |
| iCDH-2-GEMS | | SEAP | Supp Fig. 9 |
| AIR-1-GEMS | Drug-1 | SEAP | Fig. 4 |
| AIR-2-GEMS | Drug-2 | SEAP | Fig. 4 |
| MISO | | | |
| CDH-1-TF & CDH-3-TF | Drug-1 & Drug-3 | SEAP/Luciferase | Fig. 5 |
| CDH-1-GEMS & CDH-3-GEMS | Drug-2 & Drug-3 | SEAP | Fig. 5 |
| MIMO | | | |
| AIR-1-GEMS & CDH-3-TF | Drug-1 & Drug-3 | SEAP + Luciferase | Fig. 5 |
| AIR-2-GEMS & CDH-3-TF | Drug-2 & Drug-3 | SEAP + Luciferase | Fig. 5 |
| SIMO | | | |
| AIR-1-GEMS & CDH-1-TF | Drug-1 | SEAP + Luciferase | Fig. 5 |
| AIR-2-GEMS & CDH-2-TF | Drug-2 | SEAP + Luciferase | Fig. 5 |

### Methods and Materials

#### Computational design of CDH-3

The design of CDH-3 followed a similar protocol to that presented in our previous work[2], based on a side chain grafting approach. The mdm2:p53 peptide interaction was selected as a starting point, since multiple small molecule inhibitors bind to the mdm2 receptor protein and prevent binding of p53. An 8 amino acid helical fragment (**F**XXX**W**XX**L**, where F, W, and L are hotspot residues and X are the designable residues) was extracted from the helical binding motif of an mdm2-binding, p53-mimic, stapled peptide (PDB id: 5afg) and selected as the 'binding seed'. A database of monomeric protein structures obtained through X-ray crystallography was assembled, where proteins were sourced from the Protein Databank (PDB) if they met the following: (a) proteins with a global stoichiometry assigned in the PDB as monomers, (b) an amino acid sequence length between 80 and 160, (c) proteins containing helical secondary structures. The computational design protocol was executed as a script using RosettaScripts, and entailed the following steps. The MotifGraft[3] program in Rosetta attempted to graft the binding seed to all proteins in the database (*scaffold* proteins). Proteins on which the fragment was grafted to a similar backbone fragment, with a maximum $C_\alpha$ root mean square deviation of 1.0 Å, and where they maintained a steric compatibility (a clash score of maximum 5 Rosetta Energy Units or REU) with mdm2, were accepted. Once scaffolds were matched, residue positions surrounding the binding seed were designed using the Rosetta fixed backbone design[4,5], allowing amino acid mutations to residues with a positive score in the BLOSUM62 matrix[6], with respect to the amino acid identity in the wildtype protein. The restriction to mutations based on the BLOSUM62 matrix was performed to prevent mutations to the original scaffold that could affect its stability, folding pathway, or solubility. The resulting designed proteins were scored using Rosetta for the complex energy ($\Delta\Delta G$) predicted score, buried solvent accessible area and unsatisfied hydrogen bonds. Designs where Rosetta's $\Delta\Delta G$ energy was higher than -7 REU were discarded. A visual inspection of the resulting scaffolds showed that many of them had non-globular structures, with extended conformations and poor packing of the binding seed. To remove scaffolds with non-globular structures, we computed a globularity score on proteins based on a metric created by Miller et al[1], which found that the mass *M* of globular proteins correlates with the solvent accessible surface area *A* under the power law:

$$A_s = 6.3M^{0.73}$$

We thus computed a globularity score $G = A_s/6.3M^{0.73}$, and scaffolds with $G < 0.9$ were discarded from consideration. Finally, scaffolds were visually inspected, and those where a large small molecule ligand binding site was present in the original structure near the novel interface, or those where the binding seed was grafted to a terminal region of the protein, were excluded. We thus selected 3 protein scaffolds, LD4, (designed from scaffold hydroquinone flavodoxin from Desulfovibrio vulgaris with PDB id: 1AKU), LD5 (designed from scaffold with PDB id: 2IFQ) and LD6 (designed from scaffold with PDB id: 2FWE). After close inspection of LD6, residue 23 was manually mutated to Tyr, as this was the identity found in the starting stapled peptide.

#### Protein expression and purification

The genes encoding all the designs were purchased from Twist Bioscience. Genes were synthesized with N-terminal 6×His-tag and cloned into the pET11b vector using Gibson assembly (New England Biolabs, E2611S). The plasmids were transformed into XL10 gold *E.coli* for plasmid amplification and BL21(DE3) *E.coli* (Thermo Fisher) for protein expression. A single clone was picked to inoculate 500 ml of Terrific Broth (Merck Millipore, 101629) containing ampicillin (100 µg/ml). The cultures were grown at 37 °C until $OD_{600}$ reached around 0.8 and protein expression was induced with 1 mM IPTG at 20 °C. After overnight induction, cells were pelleted by centrifugation at 4,000 r.p.m. The harvested pellet (thawed on ice, if frozen after centrifugation) was resuspended in 40 ml lysis buffer (50 mM Tris, 500 mM NaCl and 5% Glycerol in pH 7.5) supplemented with 100 µg/ml PMSF (ROTH, 6376.2). The slurry was sonicated for 30 mins, centrifuged at 20,000 g for 20 mins, and the supernatant was collected. For mdm2 protein, the protein was extracted from inclusion bodies after centrifugation. The collected pellet was washed twice with 50 ml lysis buffer containing 0.05% Triton-X100 (AppliChem) and solubilized by 20 ml lysis buffer supplemented with 8 M urea (Merck, U4883) and 10 mM β-mercaptoethanol (AppliChem, A4338.0100). Resuspended inclusion bodies were dialyzed against 1 liter of 4 M guanidine hydrochloride in pH 3.5 supplemented with 10 mM β-mercaptoethanol. Next, the protein was refolded by dropwise addition with 1 liter of 10 mM Tris-HCl (pH 7.0) containing 1 mM EDTA and 10 mM β-mercaptoethanol and slowly mixed overnight at 4°C. The supernatant and the refolded solution were loaded to AKTA pure system (GE Life Science) for nickel affinity purification and the target protein was eluted by elution buffer (50 mM Tris, 500 mM NaCl, 300 mM imidazole). The eluted protein was further purified by gel filtration to separate the monodisperse population. The purified proteins were concentrated, aliquoted and stored at -80 °C.

**Compounds**

Venetoclax (>99.9%, Chemietek CT-A199), A1155463 (99.5%, Chemietek CT-A115) and NVP-CGM097 (100% optically pure, Chemitek CT-CGM097), were directly used without further purification. Venetoclax, A1155463, and NVP-CGM097 were each dissolved in DMSO as 10 mM stocks. Stocks were aliquoted and stored at -20 °C until use.

**Circular dichroism spectrum**

Folding and thermostablility of LD6 was measured using circular dichroism spectroscopy at ramping temperatures (25-90 °C). Protein samples were dissolved in phosphate saline buffer at a protein concentration around 0.2 mg/mL (20 µM). The sample was loaded into 0.1 cm path-length quartz cuvette (Hellama). The far-UV CD spectrum between 190 nm and 250 nm was recorded with a Chirascan V100 spectrometer with a band width of 2.0 nm, and scanning speed was at 20 nm/min. Response time was set to 0.125 sec and spectra were averaged from 2 individual scans.

**Size-exclusion chromatography coupled with multi-angle light scattering**

LD6 was further characterized by Size Exclusion Chromatography coupled to Light Scattering (SEC-MALS) for solution behavior, and to study dimerization and drug-induced monomerization properties. LD6 was injected at 50-100 µM into a Superdex™ 75 300/10 GL column (GE Healthcare) using a highperformance liquid chromatography system (Ultimate 3000, Thermo Scientific) at a flow rate of 0.5 ml/min. The UV spectrum at 280 nm was collected along with static light scatter signal by a MALS device (miniDAWN TREOS, Wyatt). For determining drug-induced monomerization, 50 μM mdm2 or Bcl2 were mixed with equal molarity of LD6 and LD3, respectively. Then the mixtures were treated with either 100 μM (two-fold excess) of the corresponding drugs or the same volume of DMSO to detect complex formation and forced dissociation in SEC-MALS. The light scatter signal of the sample was collected from three different angles and the results were analyzed using the ASTRA software (version 6.1, Wyatt).

**Surface plasmon resonance for assessing protein-protein binding affinity**

Surface plasmon resonance (SPR) measurements were performed on a Biacore 8K device (GE Healthcare). Drug-receptor proteins (Bcl-XL, Bcl2 and mdm2) were immobilized on a CM5 chip (GE Life Science) as a ligand with the concentration at 5 μg/ml for 120 seconds contact time in pH 4.0 (mdm2) or pH 4.5 (Bcl-XL and Bcl2) sodium acetate solutions, respectively. Serial dilutions of the analytes in HBS-EP buffer (10 mM HEPES, 150 mM NaCl, 3 mM EDTA and 0.005% Surfactant P20; GE Life Science) were flown over the immobilized chips. After each injection cycle, surface regeneration was performed using 10 mM NaOH (pH 11.95). Affinity ($K_d$) and kinetic parameters ($K_{on}$ and $K_{off}$) were obtained using a 1:1 Langmuir binding model with Biacore 8K evaluation software.

**SPR drug competition assay**

Drug $IC_{50}$s of disrupting CDH heterodimers were measured on a Biacore 8K device. 4 μM of LD3 and its variants (LD3_v1-3) or LD6 were mixed with their respective drugs. Drug concentrations ranged from 10 μM to 0.01 nM in 10x serial dilutions. The drug-binder mixtures were injected on the drug-receptor protein (Bcl-XL, Bcl2 or mdm2 as indicated) immobilized channel. Multiple-cycle injection of the protein-drug complex with different stoichiometry were performed to measure the decrease of maximal RUs (response units at 120 sec). Apparent $IC_{50}$s were obtained using the inhibition versus response fitting models in prism software (Version 8.3.0).

**Purification of mdm2 and LD6 for X-ray crystallography**

The complex of mdm2 with LD6 was prepared by mixing each of the components at equal molar ratio. After overnight incubation, the complex was purified by size exclusion chromatography using a Superdex75 16 600 (GE Healthcare) equilibrated in 10 mM Tris pH 8, 100 mM NaCl and subsequently concentrated to ~15 mg/ml (Amicon Ultra-15, MWCO 3,000). Crystals were grown using the sitting-drop vapor-diffusion method at 291K in a solution containing 1.5 M Ammonium sulfate, 0.1 M Sodium cacodylate at pH 6.5. For cryo protection, crystals were briefly swished through mother liquor containing 25% glycerol. Diffraction data were recorded with a X06DA (PXIII) beamline at the Paul Scherer Institute, Switzerland. The diffracted crystal of mdm2/LD6 belonged to space group $P4_322$. Data was integrated and processed to 2.9 Å with a high-resolution cut at I/σ=1 applied by the X-ray Detector Software (XDS)[7]. The structure was determined by molecular replacement using the Phenix Phaser module[8]. The searching of the initial phase was performed by using the mdm2 structure (PDB id: 5AFG) and the computationally designed model LD6 as a search model. Manual model building was performed using Coot[9], and automated refinement using Phenix Refine[10]. The final refinement statistics are summarized in Table S1.

**CCK8 cell viability assay**

A cell counting kit-8 (CCK-8) assay (Sigma) was used to measure the cytotoxicity of three drugs on HEK293T cells. 10,000 cells were pre-seeded into 96 well plates per well in 100 µl complete culture medium. The culture medium was changed into drug containing medium with drug concentrations of 0 µM (0.1% DMSO),1 µM, 5 µM and 10 µM. After 24 hours drug incubation, 10 µl CCK-8 solution was added to each well for 2 hours incubation under standard conditions. The absorbance at 450 nm was determined by a multiplate reader (Tecan infinite 500).

**Cell transfection and drug treatment**

HEK293T cells were maintained in DMEM medium with 10% FBS (Gibco) and Pen/Strep (Thermo Fisher). Cells were maintained and split every two days at around 80% confluence. HEK293T cells were seeded 24 hours before transfection, and transfected with the lipofectamine 3000 kit (Thermo Fisher). Plasmids were cotransfected according to the Table S7. For the drug OFF-switch experiments, drugs were added 12 hours post transfection and incubated with cells for 24 hours before the SEAP detection assay.

**SEAP detection assay**

Secreted alkaline phosphatase (SEAP) activity (U/L) in cell culture supernatants was quantified by kinetic measurements at 405 nm (1 minute/measurement for 30 cycles) of absorbance increase due to phosphatase-mediated hydrolysis of para-nitrophenyl phosphate (pNPP). 4 - 80 µL of supernatant was adjusted with water to a final volume of 80 µL, heat-inactivated (30 min at 65 °C), and mixed in a 96-well plate with 100 µL of 2 × SEAP buffer (20 mM homoarginine (FluorochemChemie), 1 mM MgCl2, 21% (v/v) diethanolamine (Sigma Aldrich, D8885), pH 9.8) and 20 µL of substrate solution containing 20 mM pNPP (Sigma Aldrich, 71768).

**Reversibility assay (Dynamic regulation)**

Cells were pre-seeded 24 hours before the transfection in 12-well plate. The ON-OFF-ON mode was grew without drug in the first 36hours and passed 1/3 of cells to the new dish with 500nM drugs supplemented for next 36hours with refreshing the drugs every 12 hours, then passed to the new dish with the removal of drugs. The OFF-ON-OFF mode was treated with 500nM drugs 12 hours post-transfection for the following 36 hours. Cells were also passed every 36 hours and cultured in the absence of drug for the hours from 36 to 72 hours, then add drugs again from 72hours. SEAP samples were taken every 12 hours from the culture supernatant.

**Design of weaker affinity variants**

The LD3 protein was computationally redesigned for decreased binding to Bcl-XL with a range of decreasing affinities. Rosetta's alanine scanning filter was used to evaluate the change in ΔΔG for the LD3:Bcl-XL complex upon mutating each of the 22 residues in the interface of LD3 to Alanine. The resulting list was then sorted by the change in ΔΔG, and three residues with positive levels of change in ΔΔG were selected: L235 (5.0 REU), D240 (4.2 REU) and E124 (1.8 REU), where higher REU values are predicted to result in greater affinity losses. The three mutations were selected to provide a 'gradient' of affinities between LD3 and Bcl-XL.

**Drug-insensitive receptor mutations predictions**

Bcl-XL and Bcl2 were redesigned for resistance to Drug-1 and Drug-2, respectively, following a computational strategy similar to one used to predict drug resistance[11,12]. Briefly, a set of residues in the receptor protein's (Bcl-XL/Bcl2) binding site was selected for redesign. From this set, a number of mutations was evaluated for binding energy to the binder protein (LD3) (positive design) or the drug (negative design). Afterwards all mutations were ranked according to the difference in energy between the positive design and negative design. Bcl-XL**:** The structure of Bcl-XL bound to Drug 1 (A-1155463, PDB id: 4QVX) was used for the negative design strategy, while the model of Bcl-XL bound to LD3 (based on the Bcl-XL:BIM BH3 structure with PDB id: 3FDL) was used for positive design. Six Bcl-XL residues in the binding site of Drug-1 (E98, R102, F105, T109, S145 and A149) were manually selected for redesign due to their proximity to drug moieties and relative distance to LD3 in the positive design structure. Each of these residues was allowed to mutate to residues with similar size/properties, restricted to a maximum of two simultaneous mutations from wildtype: E98: {E/S}, R102: {F/R/K/D/E/H}, F105:{F/L/V/I/A}; T109: {S/A/T/L/V}; S145: {S/D/E/V/A}; A149:{V/A/L/I}. The total sequence space thus consisted of 253 unique sequences. The Rosetta program was then used to redesign the positive design structure (LD3:Bcl-XL complex) and the negative design structure (Drug-1:BclxL complex) for each of the 253 sequences. A score was computed for the complex state of each sequence in each of the two states, and sequences were ranked according to the ratio of the positive design (bound) score - negative design (bound) score (Table 4, top). Five sequences iBcl-XL_v1 (T109L, A149L), iBcl-XL_v2 (A149V), iBcl-XL_v3 (R102E, F105I), iBcl-XL_v4 (R102F, T109V), and iBcl-XL_v5 (E98S, F105I) were selected from the top results. The structure of Bcl2 bound to Drug-2 (Venetoclax, PDB id: 4LVT) was used for the negative design strategy, while the model of Bcl2 bound to LD3 (PDB id: 6IWB) was used for positive design. Five Bcl2 residues in the binding site of Drug-2 (A100, D103, V148, V156 and Y202) were manually selected for redesign due to their closeness to drug moieties and relative distance to LD3 in the positive design structure. Each of these residues was allowed to mutate to amino acids with similar size/properties, restricted to a maximum of two simultaneous mutations from wildtype: A100: {A/S/T/V}, D103: {D/N/E/Q/S}, V148: {V/I/L/M/T}, V156: {V/I/L/M/T} and Y202: {Y/W/F/H/R/K/Q/E}. The total sequence space thus consisted of 251 unique sequences. The Rosetta program was then used to redesign the positive structure (PDB id: 6IWB) and the negative design structure (PDB id: 4LVT) for each of the 251 sequences. A score was computed for the complex state of each sequence in each of the two states, and sequences were ranked according to the ratio of positive design score/negative design score (Table 5, top). Three sequences iBcl2_v1(156I, 202H), iBcl2_v2(103N, 202H) and iBcl2_v3(100T_103S) were selected from the top results. Three additional mutations were enriched in the top designs, and therefore we designed a further version, iBcl2_v4 (100V_103N_202H).

**Statistics**

Apparent $IC_{50}$s of SPR drug competition assay were calculated using three-parameter nonlinear regression in GraphPad Prism (Version 8.3.0). Representative data of cellular assays are presented as individual values and mean values (bars). n = 3 refers to biological replicates. All $IC_{50}$/$EC_{50}$ values of cellular assays (CDH-TF, CDH-GEMS and AIR-GEMS) reported were calculated using four-parameter nonlinear regression ± s.d.
